# The claudin-like molecule CLC-3 regulates neuromuscular function in *Caenorhabditis elegans* by modulating cholinergic signalling

**DOI:** 10.1101/2025.08.18.670992

**Authors:** Aishwarya Ahuja, Anusha Rastogi, Haowen Liu, Akankshya Sahu, Harinika Kumar, Jagannath Jayaraj, Zhitao Hu, Kavita Babu

## Abstract

Cell adhesion molecules (CAMs play important roles in neurons, contributing to nervous system development, synapse formation, and activity-dependent plasticity. Claudins, the cell-adhesion molecules known for their roles at tight junctions in epithelial and endothelial cells, remain underexplored in neurons, particularly in vertebrates. In contrast, emerging studies in *Caenorhabditis elegans* have begun to reveal neuronal functions of claudin-like proteins. However, a systematic analysis of their neuronal expression has not been performed. We conducted a transcriptional reporter screen of all claudin-like genes in *C. elegans* and identified several candidates with previously unreported neuronal expression, highlighting a broader role of this family in the nervous system. One candidate, *clc-3*, showed robust expression in head, tail, and ventral cord neurons, with no detectable expression in non-neuronal tissues. Functional analyses of *clc-3* mutants revealed increased body-bend amplitude and elevated evoked postsynaptic currents at the cholinergic neuromuscular synapses. Imaging and molecular interaction studies demonstrated that CLC-3 localises to the presynaptic membranes in cholinergic neurons, where it interacts with the actin-binding protein NAB-1 and regulates cholinergic signalling. This presynaptic role of CLC-3 likely contributes to the regulation of sinusoidal movement in *C. elegans*. Our findings identify CLC-3 as a neuronally expressed claudin that regulates motor system output by influencing synaptic vesicle organization and illustrate how changes in synaptic organization are coupled to whole-animal behaviour.

## Introduction

Cell adhesion molecules (CAMs) are present at the junction of many cells, including interneuronal synapses and neuron-glia contacts, and are key to nervous system development and maintenance (*1*). In *Caenorhabditis elegans*, like in other organisms, CAMs regulate various aspects of neural function, including axon outgrowth, synapse recognition and formation, synaptic plasticity, neuronal positioning, and neural circuit maintenance (*2–16*). Consequently, disruptions in the function of these molecules can lead to impairments in nervous system function, often reflected as alterations in the animal’s behaviour. For instance, *C. elegans* mutants lacking a cell adhesion molecule neurexin exhibit defects in their response to gentle touch in both anterior and posterior parts of the body when compared to wild-type (WT) animals, likely due to defects in synaptic signalling (*17*). Additionally, mutations in other CAMs, such as *rig-3*, result in altered reversal frequency during locomotion, while *ncam-1* mutants show defects in long-term associative memory. (*18, 19*). Despite the remarkable diversity of CAMs functioning in neurons, one particular family of transmembrane proteins remains relatively overlooked in this context– the claudins.

Claudins are tetraspan membrane proteins that serve as core structural components of tight junction complexes in epithelial and endothelial cells of vertebrates, where their roles are primarily adhesive (*20–25*). Within the nervous system, these proteins play a critical role in sealing spaces between brain microvascular endothelial cells, thereby contributing to the formation and maintenance of the blood-brain barrier (*26–32*). While cell-adhesion roles of claudins at junctional complexes have been well characterised, studies that examine their expression and function within neurons remain limited. Nonetheless, a few reports suggest neuronal expression of claudins in vertebrate systems. For instance, a study has observed expression of claudins 2, 5 and 11 in pyramidal neurons of the frontal cortex in post-mortem human brain tissue (*33*). In mice, Claudin-12 expression has been reported in neuronal populations of brain regions including the cortex, hippocampus, and cerebellum (*34*).

In contrast to the limited data available on claudin expression within vertebrate neurons, studies on an invertebrate model system, *C. elegans*, have begun to shed light on neuronal functions of claudins. Although classical tight junctions are absent in *C. elegans*, its genome encodes at least 17 claudin-like molecules that share structural and functional homology with their vertebrate counterparts and are thought to regulate cell adhesion and paracellular permeability (*35–37*). The simplicity of its nervous system, combined with the ease of genetic manipulations, makes *C. elegans* a powerful model system to study previously uncharacterised functions of proteins (*38–41*). Indeed, these advantages have facilitated the discovery of neuronal and synaptic roles for at least three claudin-like molecules in *C. elegans* (*42*): 1) HIC-1 is implicated in regulating postsynaptic acetylcholine (ACh) receptor levels via Wnt signalling at cholinergic synapses (*43*); 2) NSY-4 mediates lateral signalling between the AWC neuron pair (*44*); and 3) HPO-30 contributes to dendritic branching in the PVD neuron in addition to functioning at the neuromuscular junction to maintain levamisole sensitive ACh receptor levels (*45–47*).

Despite such important insights, no comprehensive analysis has yet been performed to determine the neuronal expression of the entire claudin set in *C. elegans*. To address this gap, we performed a reporter-based screen utilizing fluorescently tagged transcriptional constructs to assess neuronal expression of *C. elegans* claudins. We identified nine claudins that show neuronal expression, with one candidate, *clc-3*, showing strong and exclusive expression in neuron populations including the head, tail and ventral cord neurons. Behavioural analyses of *clc-3* mutants revealed a defect in the sinusoidal body-bend amplitude, suggesting a role for this claudin in regulating motor system output.

Further investigations into the function of CLC-3 revealed a presynaptic role of the claudin in *C. elegans* cholinergic neurons, where it regulates width of synaptobrevin puncta, a neurotransmitter vesicle-associated protein, along the dorsal nerve cord (DNC) via interactions with an actin-binding protein NAB-1. Our findings reveal a fine-tuning function of CLC-3 in regulating presynaptic functions and motor-circuit dynamics, and demonstrate how subtle changes in synaptic organization can influence quantifiable behavioural outputs. This work provides a direct link between synaptic signalling and locomotor behaviour and adds to the growing evidence that claudins perform roles beyond their traditional functions at tight junctions.

## Materials and Methods

### Strain maintenance

All *C. elegans* strains were maintained on nematode growth medium (NGM) plates seeded with *E. coli* OP50 bacteria, under standard laboratory conditions at 22°C, following well-established protocols described in literature (*48*). Single mutants obtained from the Caenorhabditis Genetics Center (CGC) were outcrossed 3-4 times with the wild-type (WT) N2 Bristol strain which served as the standard WT control across all experimental analyses. Synchronised populations of young adult *C. elegans* were obtained using standard synchronization protocols (*49*). A complete list of all the strains used in this study is provided in Supplementary Table 1, and primers used for genotyping are listed in Supplementary Table 2. The *nuIs152, nuIs159* (*50*), and *nuIs299* (*6*) strains were gifts from Josh Kaplan’s lab.

### Molecular cloning and transgenic generation

Promoters for all transcriptional reporter constructs were cloned into the multiple cloning site (MCS) of the plasmid pPD95.75 using Gibson assembly method. Rescue constructs for CLC-3 expression were generated by cloning the entire genomic region of CLC-3 under 2-3kb promoter sequence. For the variant lacking PDZ-binding motif, the genomic sequence lacking the last seven amino acids was used. In both cases, the promoter and gene sequences were cloned upstream of and in frame with a fluorescent reporter (GFP or mCherry), separated by a self-cleaving T2A sequence. The sequences of all promoters and genes were extracted from WormBase. The sequences for the split YFP sequence used in BiFC experiments were amplified from Addgene plasmids pBiFc-VN173 (#22010) and pBiFc-VC155 (#22011). All constructs were microinjected in *C. elegans* gonads to generate transgenic lines as previously described (*51, 52*). The list of all plasmids used and generated in this study is provided in Supplementary Table 3. List of primers used for cloning is provided in Supplementary Table 4.

### Locomotion assays

Locomotion assays were performed with age-synchronised young adults across all strains. Briefly, a single animal was picked using an eyelash pick and placed onto a 90mm NGM plate lacking food, allowing it to crawl freely to get rid of residual food sticking to the *C. elegans* body. Subsequently, the animal was transferred to a no-cholesterol, 90 mm NGM assay plate without food and its movement was recorded for 5 minutes at room temperature (maintained at 22°C) using the WormLab imaging system. The average body-bend amplitude and speed of locomotion were analysed using the WormLab software. The number of *C. elegans* recorded for each genotype was ≥ 20.

### Aldicarb Assays

Aldicarb assays were performed as described previously with modifications (*53*). Plates containing 3 mM Aldicarb (BIOELSA) were prepared one day before the assay and allowed to dry at room temperature. Young adult animals (20–25) were picked and placed on aldicarb plates. The animals were scored for paralysis by gently prodding them thrice on the head and thrice on the tail after every 20 mins with an eyelash pick. The *C. elegans* were characterised as paralysed if they showed no movement in response to touch. Most assays were performed in triplicate, with the experimenter being blind to the genotype of the animals. Due to the unavailability of the chemical aldicarb towards the end of the study, certain assays, specified in the Results section, were performed either twice or as a single replicate. For simplicity, the percentage of animals that were paralyzed at the 160 or 180 min time point was plotted.

### Electrophysiology Recordings

Electrophysiology recordings were performed on dissected *C. elegans* as previously described (*16, 54*). The *C. elegans* were superfused in an extracellular solution containing 127 mm NaCl, 5 mm KCl, 26 mm NaHCO_3_, 1.25 mm NaH_2_PO_4_, 20 mm glucose, 1 mm CaCl_2_, and 4 mm MgCl_2_, bubbled with 5% CO_2_/95% O_2_ at 20°C. Whole-cell recordings were performed at −60 mV (reversal potential of GABAA receptors) for mEPSCs and 0 mV (reversal potential of ACh receptors) for mIPSCs. The internal solution contains 105 mm CsCH_3_SO_3_, 10 mm CsCl, 15 mm CsF, 4 mm MgCl_2_, 5 mm EGTA, 0.25 mm CaCl_2_, 10 mm HEPES, and 4 mm Na_2_ATP, adjusted to pH 7.2 using CsOH. Stimulus-evoked EPSCs were stimulated by placing a borosilicate pipette (51 μm) near the ventral nerve cord (one muscle distance from the recording pipette) and applying a 0.4 ms, 85 μA square pulse (WPI). To measure ACh -activated currents, a puffing pipette (5–10 μm open size) containing ACh was placed at the end of the patched muscle, and pressure was applied via Picospritzer (Parker).

### Microscopy and imaging

Images were obtained using an assembled epifluorescence scope. To prepare *C. elegans* for imaging experiments, animals in the young adult stage were immobilised using 30 mg/ml 2, 3-butanedione monoxime (BDM) on 1% agarose pads. The analysis of images was done using the ImageJ software. Average fluorescence intensity, and puncta density were quantified and plotted on graphs using the Prism software (GraphPad).

### BiFC experiments

To perform the BiFC assay, VN173 Fragment (N-terminus) of split YFP was cloned downstream to Neurabin cDNA, and VC155 Fragment (C-terminus) of split YFP was tagged to the C-terminus of CLC-3 genomic DNA (*55*). For the control experiment, CLC-3ΔPDZ(7aa)::VC155 was generated. The YFP fragments were connected to their respective proteins via a flexible linker. All the constructs for BiFC experiments were cloned under the *unc-17* promoter to drive the expression in the cholinergic neurons.

### Statistical analysis

The data were plotted using GraphPad Prism 10, and statistical analyses were performed using either Student’s *t*-test or one-way ANOVA with multiple comparisons, as specified in figure legends. A p-value of ≤ 0.05 was considered statistically significant. Data are presented as either mean ± SEM or mean ± SD, as indicated in the respective figure legends.

## Results

### *Caenorhabditis elegans* claudin-like proteins are expressed in neurons

The *C. elegans* genome encodes at least 17 claudin-like proteins that exhibit both structural and functional homology to their vertebrate counterparts ((*37, 45*) and WormBase)). While a subset of these claudins, including HPO-30, HIC-1 and NSY-4, has been previously shown to be expressed and functionally active in neurons, other candidates display an expression pattern more typical of classical claudins (*43–45, 47*). For example, CLC-1 is expressed in epithelial cells of the pharyngeal region, CLC-2 localises to the seam cells of the hypodermis and VAB-9 is expressed broadly in all epithelial cells as well as within the nerve ring (*36, 56*). Despite these insights, many members of the *C. elegans* claudin family remain uncharacterised, and no comprehensive analysis has yet been performed to systematically define the neuronal expression of the entire claudin repertoire.

To investigate whether additional members of the *C. elegans* claudin family are expressed in neurons, we conducted a transcriptional reporter screen, where the promoter for each claudin, a 2-3 kb sequence upstream of the start codon, was tagged to the green fluorescent protein (GFP) and microinjected into the *C. elegans* gonads. The resulting transgenic lines were examined to determine the expression patterns of the respective claudin genes (Figure 1A, 1B).

**Figure 1:**
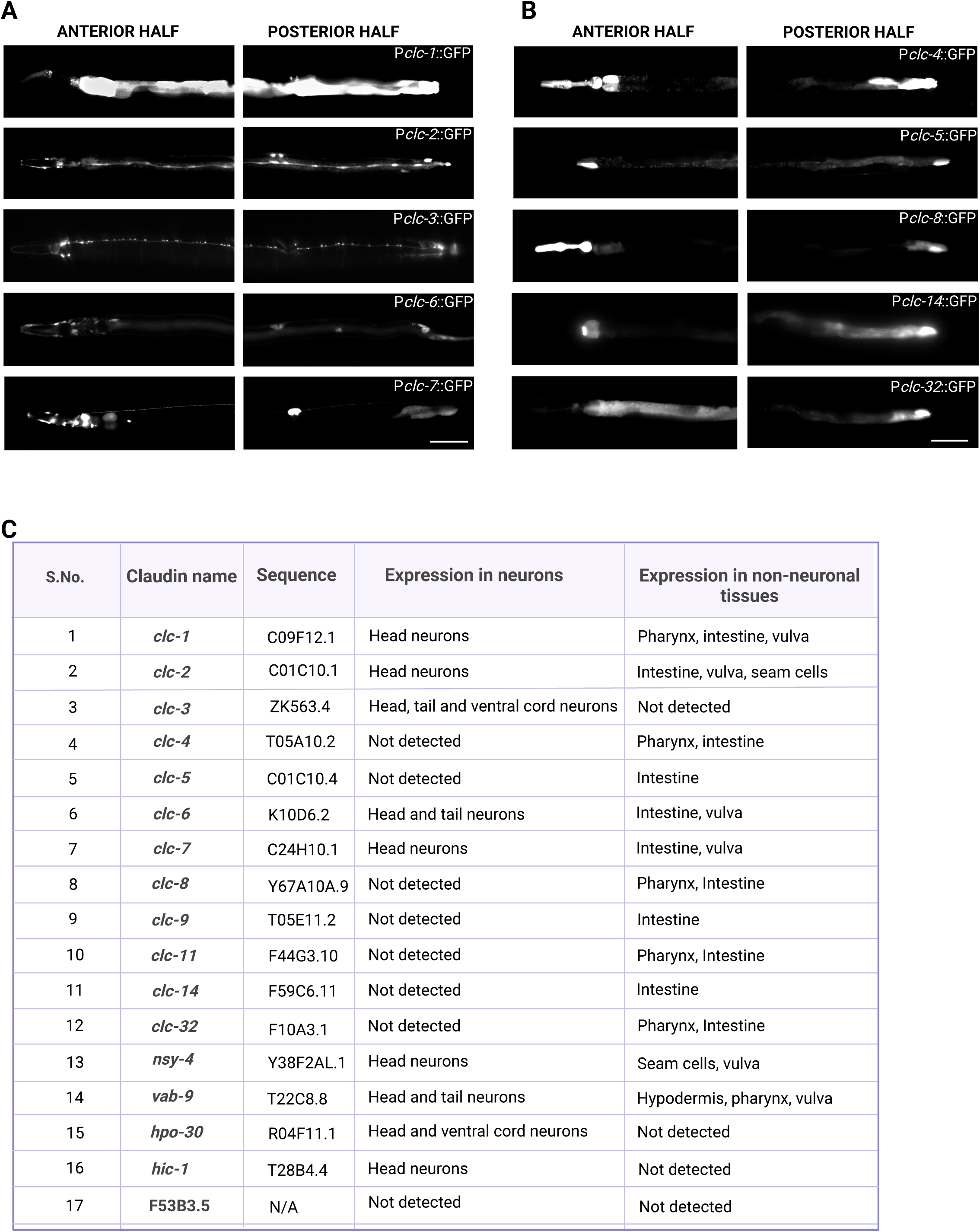
*C. elegans* claudin-like proteins are expressed in neurons. (A) Representative images showing transcriptional reporter expression of claudin-like genes in wild-type (WT) *C. elegans*. Reporters for *clc-1*, *clc-2*, *clc-6*, and *clc-7* are expressed in both neuronal and non-neuronal tissues, while *clc-3* shows robust and specific expression in neuronal tissues, including head, tail, and ventral cord neurons. Scale bar = 100 μm (B) Expression of transcriptional reporters for *clc-4*, *clc-5*, *clc-8*, *clc-14*, and *clc-32* is limited to non-neuronal tissues and is not detected in neurons. Scale bar = 100 μm (C) Table summarizing the tissue-specific expression patterns of transcriptional reporters for claudin-like genes.

We found that as many as nine claudin candidates exhibited significant neuronal expression (Summarised in the table in Figure 1C). Notably, both *clc-1* and *clc-2*, which were previously uncharacterised for neuronal expression, were found to be expressed in head neurons of *C. elegans* (Figure 1A). Additionally, claudins *clc-6* and *clc-7* displayed expression in neuronal as well as non-neuronal tissues. While both *clc-6* and *clc-7* were expressed in multiple head neurons and the vulva region, *clc-6* also showed expression in the intestine (Figure 1A). Interestingly, claudin *clc-3* exhibited robust and largely exclusive expression in neurons including head neurons, tail neurons and ventral cord neurons (Figure 1A). This observation is consistent with the single-cell RNA sequencing data from the CeNGEN project, which indicates that the *clc-3* transcript is enriched in head neurons such as AWB, AIA, and AIZ; ventral cord motor neurons, including VC4, VC5, VA12, DA9, VC, and AS; and tail neurons such as LUA and PDB (*57, 58*).

The expression patterns of *nsy-4*, *hic-1*, *hpo-30* and *vab-9* in *C. elegans* neurons largely recapitulated previously explored expression profiles, although *vab-9* also showed potential expression in a few tail neurons (Figure S1A). In contrast, the expression of the remaining claudin candidates was limited to non-neuronal tissues, primarily the pharynx and intestine. Among these, claudins *clc-4*, *clc-8*, *clc-11* and *clc-32* were expressed in both pharyngeal and intestinal regions, whereas others, including *clc-5*, *clc-14* and *clc-9* were expressed only in the intestine (Figures1B, S1A and S1B). No expression was detected in neuronal or non-neuronal tissues for the claudin *F53B3.5* (table in Figure 1C).

Based on these findings, *clc-3* was chosen as the candidate to study the neuronal functions of claudins, as it stood out due to its strong and predominantly neuronal expression pattern.

### Mutants in *clc-3* exhibit deeper body-bend amplitudes

*C. elegans* moves on the surface of agar plates in an undulatory fashion. The ventral cord in these animals harbours motor neurons that synapse onto the dorsal and ventral body wall muscles, regulating their excitation and inhibition in a coordinated manner to ensure alternating muscle contraction (*59*). This alternating contraction generates the characteristic sinusoidal wave of movement. Disruptions in the excitation-inhibition balance have been associated with abnormalities in sinusoidal locomotion. For example, increased synaptic activity and consequent muscle excitability, driven by enhanced activity of levamisole-sensitive ACh receptors, has been linked with an elevated sinusoidal wave amplitude (*60*).

Since *clc-3* showed strong expression in the ventral cord neurons, we obtained *clc-3* mutants carrying a 1200 bp deletion in the genomic region from CGC and examined them for locomotory defects related to the sinusoidal motion of *C. elegans* (Illustrated in Figure 2A). The movement of well-fed young adult animals on a no-food NGM plate was recorded for 5 minutes using the WormLab Tracker system (Assay Illustrated in Figure S2A). We observed that while *clc-3* mutants exhibited no significant difference in their locomotion speed when compared to wild-type (WT) *C. elegans* (Figure S2B), the amplitude of their sinusoidal wave (Illustrated in Figure 2B) was markedly increased (Figure 2C). To ensure that this phenotype was in fact mediated by CLC-3, we expressed CLC-3 under its endogenous promoter in the mutant background, which resulted in a complete rescue of the amplitude defect (Figure 2C and worm tracks are indicated in Figure S2C). These findings suggest that CLC-3 plays a role in regulating the sinusoidal wave amplitude during locomotion.

**Figure 2:**
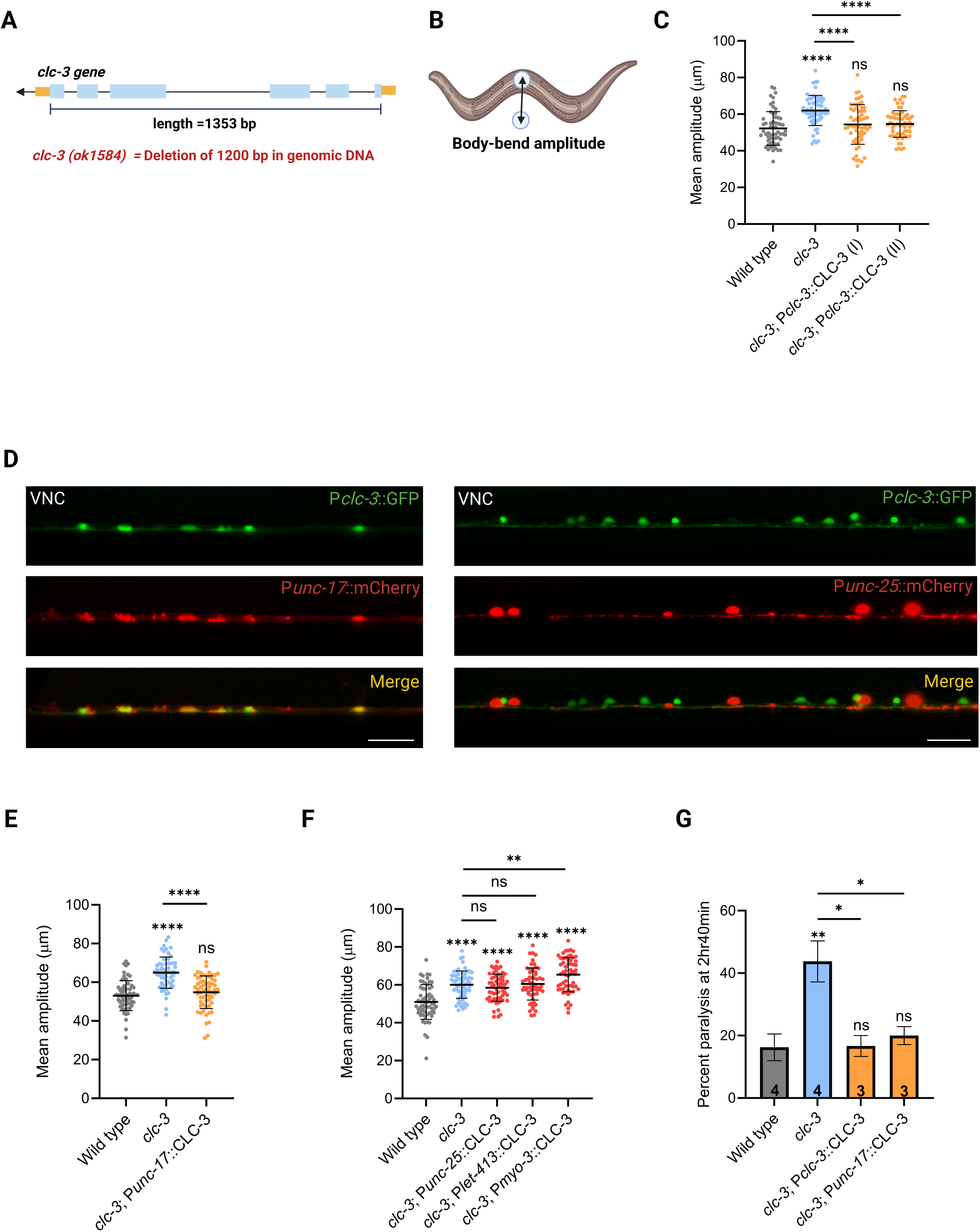
Mutants in *clc-3* exhibit deeper body-bend amplitude. (A) Illustration of the *clc-3* genomic locus showing exons (blue) interspersed with introns. Untranslated regions (UTRs) are highlighted in yellow. (B) Schematic illustrating body-bend amplitude measurement during *C. elegans* sinusoidal movement. (C) Quantification of mean body-bend amplitude recorded over five minutes of movement. *clc-3* mutants exhibit increased body-bend amplitude when compared to WT animals. This phenotype is fully rescued by expressing CLC-3 under its endogenous promoter, as observed in two independent rescue lines (I and II). Each dot represents the average body-bend amplitude over a 2.5-minute interval. For each animal, two data points are shown, corresponding to the first and second halves of the 5-minute recording. Number of trials: *N*=3; number of *C. elegans* used per trial: *n* =10. Data are mean ± SD, \*\*\*\**p* < 0.0001. (D) The *clc-3* transcriptional reporter is expressed in cholinergic ventral cord neurons (marked by P*unc17*::mCherry; left), but not GABAergic ventral cord neurons (marked by P*unc25*::mCherry; right). Scale Bar = 20 μm (E) The increased body-bend amplitude observed in *clc-3* mutants is fully rescued by expressing CLC-3 under the cholinergic neuron-specific promoter P*unc-17*. *N*=3; *n*=10. Data are mean ± SD, \*\*\*\**p* < 0.0001. (F) Targeted expression of CLC-3 under GABAergic (P*unc-25*), epithelial (P*let-413*), or body-wall muscle (P*myo-3*) promoters fails to rescue the elevated body-bend amplitude observed in *clc-3* mutants. *N*=3; *n*=10. Data are mean ± SD, \*\*\*\**p* < 0.0001. (G) Bar graph showing the percentage of animals paralyzed at the 160-minute time point following aldicarb exposure. *clc-3* mutants display hypersensitivity to aldicarb, which is rescued by expressing CLC-3 under either its endogenous or cholinergic neuron-specific promoter. In all aldicarb assays, the number at the base of each bar indicates the number of independent trials performed, with *n*=20 *C. elegans* per trial. Data are mean ± SEM, \*\**p*=0.0045. For all graphs in this figure One-way ANOVA followed by Tukey’s post-hoc test was performed to compare results.

To determine whether CLC-3 functions in neurons, we were interested in performing neuron-specific rescue experiments by expressing CLC-3 in defined neuronal subsets. As the ventral cord harbours both cholinergic and GABAergic neurons, we first identified the neuronal population expressing *clc-3*. Co-localization experiments with cholinergic and GABAergic specific markers revealed that *clc-3* is expressed in cholinergic neurons but not GABAergic neurons (Figure 2D). Consistent with this, expression of CLC-3 under a cholinergic promoter fully rescued the amplitude defect, whereas expression under a GABAergic promoter failed to restore normal amplitude (Figures 2E, F and track images indicated in S2C). While *clc-3* did not exhibit detectable expression in body wall muscles or epithelial cells in the transcriptional reporter screen, we sought to rule out contributions from these tissues by expressing CLC-3 under muscle and epithelial promoters in the mutant background. Neither expression rescued the amplitude defect, further supporting the hypothesis that CLC-3 acts primarily via cholinergic neurons to regulate sinusoidal wave amplitude (Figure 2F).

To investigate if changes in synaptic signalling underlie the amplitude phenotype observed in *clc-3* mutants, we performed an aldicarb assay. In this assay, *C. elegans* are exposed to aldicarb, an inhibitor of the enzyme responsible for breaking down the neurotransmitter ACh (*53*). We saw that *clc-3* mutants were hypersensitive to aldicarb, which may reflect either increased excitatory signalling or decreased inhibitory input at the synapse (Figure 2G). This hypersensitivity was fully rescued when CLC-3 was expressed under its endogenous promoter or a cholinergic neuron-specific promoter (Figure 2G and Figure S2D).

Based on these findings, we hypothesised that CLC-3 regulates body-bend amplitude in *C. elegans* by modulating synaptic signalling in the cholinergic neurons.

### Mutants in *clc-3* display an increase in evoked currents at the neuromuscular junction

We used electrophysiology recordings to probe further into synaptic signalling in *clc-3* mutants. Recordings from the body-wall muscles in these mutants revealed that the spontaneous excitatory and inhibitory postsynaptic currents (EPSCs and IPSCs) were unaffected, indicating that the baseline neurotransmitter release remains normal in absence of *clc-3* (Figures 3A and B). Additionally, the response of the body wall muscles to exogenous application of the neurotransmitter ACh was comparable between mutants and WT animals, suggesting that the postsynaptic receptor sensitivity in mutants is unaltered (Figure S3A).

**Figure 3:**
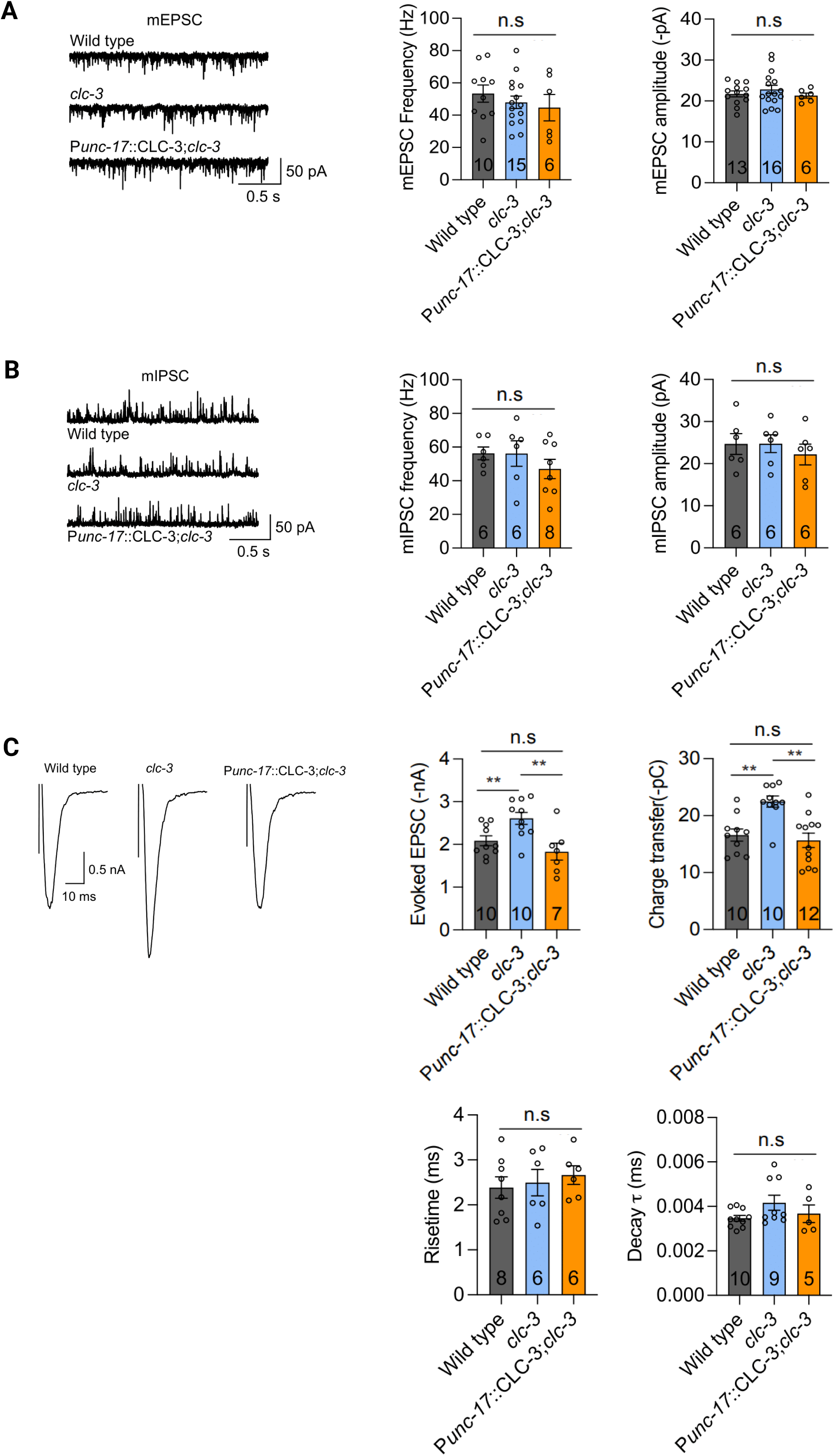
Mutants in *clc-3* display an increase in evoked currents at the neuromuscular junction. (A) Representative traces of whole-cell recordings from body wall muscles showing miniature excitatory postsynaptic currents (mEPSCs). Quantification of mEPSC frequency and amplitude, shown in the accompanying bar graphs, indicates no significant difference between *clc-3* mutants and WT *C. elegans*. (B) Representative traces showing miniature inhibitory postsynaptic currents (mIPSC). Quantification of mIPSC frequency and amplitude indicates that these parameters are unaltered in *clc-3* mutants. (C) Trace of stimulus-evoked response recorded from the body wall muscles following electrical stimulation of the ventral cord. Quantification of evoked current amplitude and total charge transfer reveals a significant increase in these parameters in *clc-3* mutants. These changes are rescued upon expression of CLC-3 under a cholinergic neuron-specific promoter. Numbers at the base of the bars indicate the number of *C. elegans* used for recordings. Data are mean ± SEM.

However, when the ventral cord in *clc-3* mutants was stimulated with an electrical impulse, the resulting postsynaptic currents in the body wall muscles were significantly elevated in comparison to those seen in WT animals, as indicated by an increase in amplitude and charge transfer of evoked EPSCs (Figure 3C). Importantly, this increase in evoked release was rescued when CLC-3 was expressed under the cholinergic promoter in the mutant background (Figure 3C).

These findings suggest that CLC-3 acts in cholinergic neurons to regulate action-potential derived neurotransmitter release, without affecting spontaneous synaptic transmission or postsynaptic receptor function.

### Synaptic vesicle distribution is altered in *clc-3* mutants

To investigate the molecular basis of elevated evoked currents in *clc-3* mutants, we performed imaging analyses using multiple fluorescently tagged markers, including those for synaptic proteins and body-wall muscles. We first found that the expression of CLC-3 overlapped with that of the synaptic vesicle-associated protein SNB-1 along the DNC in WT *C. elegans* (Figure 4A). To examine the neurotransmitter vesicles in *clc-3* mutants, we visualised GFP-tagged SNB-1 in cholinergic neurons of these mutants. We found that the overall abundance of SNB-1 puncta in mutants appeared largely normal, as indicated by normal values for average fluorescent intensity and punctal density (Figure 4B and Figure S4A). However, there was a marked increase in the width of the puncta, pointing towards a defect in vesicle localization, with SNB-1 puncta appearing broader and more aggregated in *clc-3* mutants compared to WT animals (Figure 4B). Notably, this defect in punctal width was fully rescued by specifically expressing CLC-3 in cholinergic neurons (Figure 4B).

**Figure 4:**
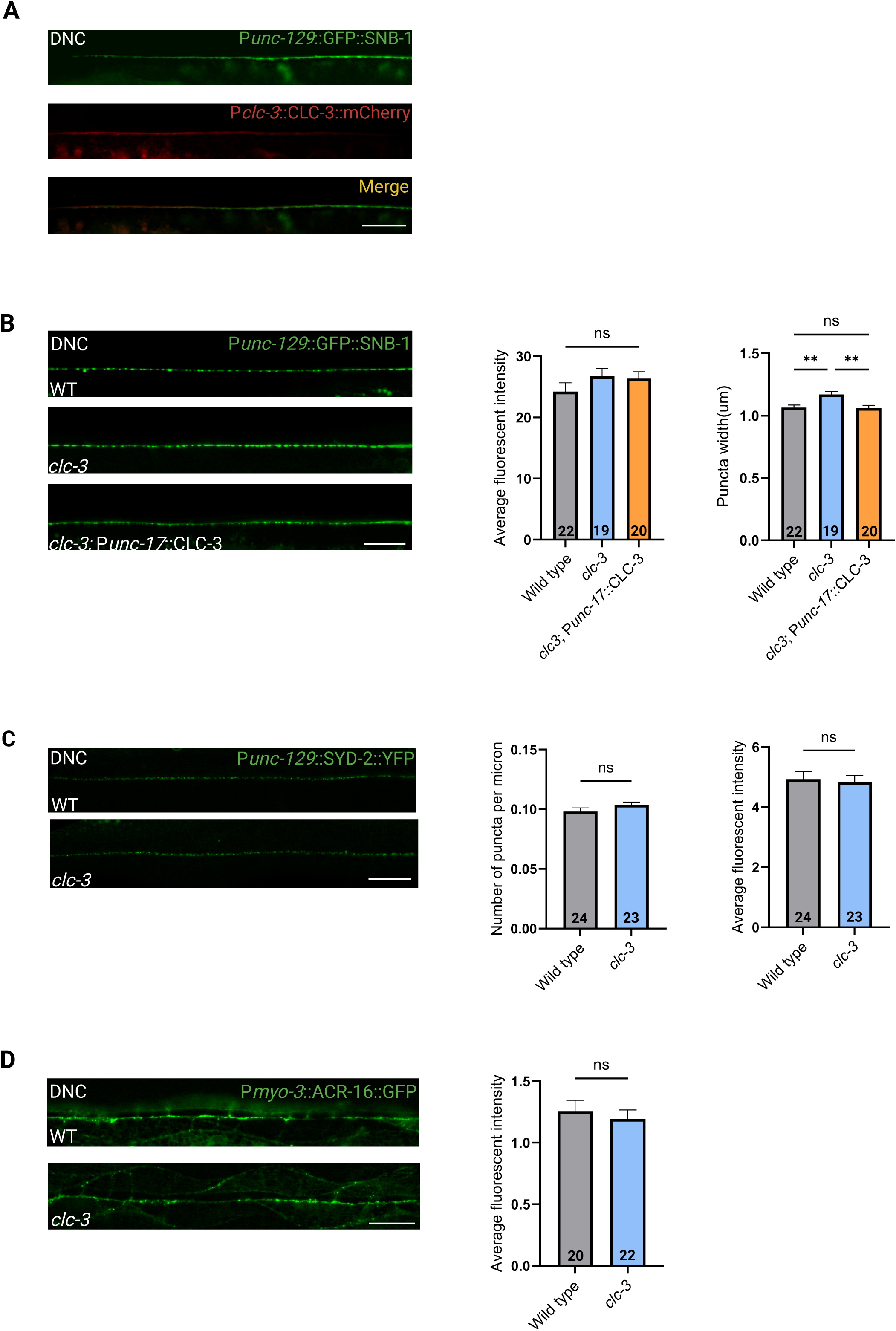
Synaptic vesicle distribution is altered in *clc-3* mutants. (A) Representative images showing colocalization of SNB-1 and CLC-3-along the DNC at cholinergic synapses. The expression of CLC-3 overlaps with SNB-1 at presynaptic regions along the DNC. Scale bar = 20 μm (B) Representative images showing the distribution of SNB-1 puncta at the cholinergic synapses along the DNC. Scale bar = 20 μm.. The SNB-1 puncta appear to be more closely clustered in *clc-3* mutants, with increased puncta width (right) but no change in the average fluorescent intensity (left), as shown in the accompanying bar graphs. The increase in puncta-width is rescued when CLC-3 is expressed under a cholinergic neuron-specific promoter. Data are mean ± SEM, \*\**p*=0.001. (C) Representative images showing SYD-2 expression at presynaptic sites in cholinergic neurons along the DNC of WT and *clc-3* mutant animals. Scale bar = 20 μm. Quantification of puncta density and average fluorescent intensity indicates normal expression of the active zone marker SYD-2 in *clc-3* mutants. (D) Representative images showing ACR-16 expression at the postsynaptic sites along the DNC in WT and *clc-3* mutant animals. Scale bar = 20 μm. ACR-16 levels are unaltered in *clc-3* mutants, as indicated by comparable average fluorescent intensity in the accompanying bar graph.. Numbers at the base of the bars indicate the number of *C. elegans* used for imaging. Statistical analyses were performed using one-way ANOVA followed by Tukey’s post-hoc test for (B) and unpaired Student’s *t* test for (C) and (D).

We also attempted to determine if the elevated wave amplitude in *clc-3* mutants could be attributed to broader defects in synaptic morphology or development. To assess the presynaptic structures, we visualised the expression of YFP tagged SYD-2, an active zone marker, in cholinergic neurons, and found no significant changes in synapse morphology between *clc-3* mutants and WT animals (Figure 4C). Similarly, the expression of the postsynaptic ACh receptor ACR-16 appeared unchanged in *clc-3* mutant animals, suggesting that postsynaptic organization in the mutants is intact (Figure 4D). These observations are consistent with our electrophysiological data, which showed normal spontaneous postsynaptic currents and unaltered responses to exogenous ACh in mutants.

To rule out defects in the development of cholinergic neurons, we used a transcriptional reporter expressing (mCherry) under a cholinergic promoter. The neuronal architecture of cholinergic neurons in *clc-3* mutants was preserved and did not show any developmental defects (Figure S4B). Additionally, the synaptic contacts between neurons and the body wall muscles, representing the NMJ, appeared normal in *clc-3* mutants, indicating proper NMJ development in the mutants (Figure S4C). We also examined whether structural defects in the body wall muscles contributed to the heightened wave amplitude in mutants. To this end, we visualised the expression of GFP-tagged MYO-3, which encodes a myosin heavy chain, an integral component of the contractile machinery (Figure S4D). We found that the muscle organization in *clc-3* mutants was indistinguishable from that in WT animals, ruling out muscle abnormalities as a contributing factor.

Thus, our data indicate that synaptic and muscle development in *clc-3* mutants is largely normal. The sole detectable abnormality was the increased width of SNB-1 puncta, indicating towards a defect in the organization of the vesicle pool at the presynapse in these mutants. This structural abnormality may underlie the enhanced levels of evoked release and body-bend amplitude observed in *clc-3* mutants.

### The PDZ-binding motif of CLC-3 is required for regulating body bend amplitude

To delve deeper into the mechanism underlying CLC-3 function in cholinergic neurons, we examined the structural features of CLC-3 that might contribute to the body-bend amplitude defect in mutants. Claudins, both in vertebrates as well as *C. elegans,* exhibit a conserved structural architecture consisting of two extracellular loops, a short intracellular loop, and intracellular N-and C-terminal domains (*21, 35*). A characteristic feature of many claudins is the presence of a PDZ-binding motif at the C-terminus that facilitates interactions between claudins and PDZ-domain containing scaffolding proteins, thereby influencing downstream signalling pathways (*61*). CLC-3 also harbours a PDZ-binding motif at its C-terminus, and given its presynaptic role in cholinergic neurons, we hypothesised that this motif may be important for mediating intracellular protein-protein interactions at the presynapse, potentially contributing to the body-bend amplitude defect in *clc-3* mutants.

To test this, we generated transgenic lines expressing a version of CLC-3 protein lacking the PDZ-binding motif, driven either by its endogenous promoter or a cholinergic neuron-specific promoter, in the mutant background (Illustrated in Figure 5A). We found that both constructs failed to rescue the body-bend amplitude defect in *clc-3* mutants (Figure 5B). Additionally, neither transgene could rescue the hypersensitivity of the mutants to aldicarb (Figure 5C, S5A and S5B). However, these experiments were performed toward the end of the study, and due to the unavailability of the chemical aldicarb in India, the assay could only be repeated twice for the endogenous promoter construct and once for the cholinergic promoter. While the limited data set precludes robust statistical analysis and constrains our ability to draw definitive conclusions regarding the rescue of aldicarb hypersensitivity in the absence of PDZ-binding motif, imaging analyses of GFP tagged SNB-1 in transgenic animals expressing CLC-3 lacking the PDZ-binding motif under the cholinergic promoter revealed a failure to rescue the SNB-1 puncta aggregation defects observed in *clc-3* mutants (Figure 5D), while punctal density appeared unaffected in all cases (Figure S5C). Based on these findings, it is reasonable to infer that the PDZ-binding domain likely contributes to synaptobrevin punctal width at the presynapse and plays a role in the altered body-bend amplitude phenotype.

**Figure 5:**
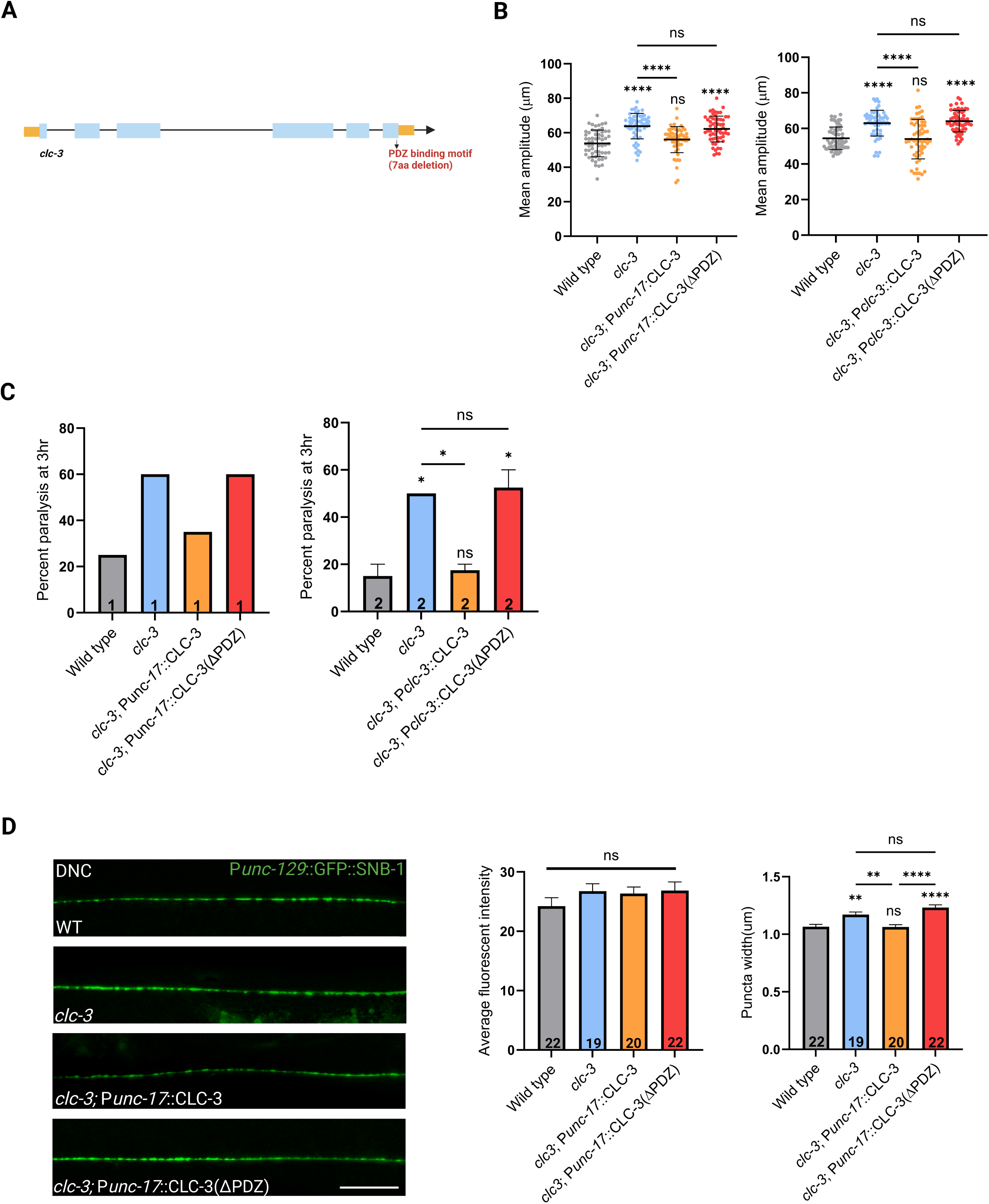
The PDZ-binding motif of CLC-3 is required for regulating body bend amplitude. (A) Schematic of the *clc-3* with a deletion of the PDZ-binding motif at the C-terminus. (B) Deletion of the PDZ-binding motif prevents the rescue of body-bend amplitude defect in *clc-3* mutants. Bar graphs show quantification of the mean body-bend amplitude in rescue lines expressing CLC-3 lacking the PDZ bm, driven either by the endogenous promoter (left) or a cholinergic neuron-specific promoter (right) in the mutant background. *N*=3; *n*=10. Data are mean ± SD, \*\*\*\**p*<0.0001. (C) Bar graphs showing the percentage of animals paralyzed at the 180-minute time point following aldicarb exposure. Expression of CLC-3 lacking the PDZ bm, either under the endogenous promoter (left) or a cholinergic neuron-specific promoter (right), fails to rescue the hypersensitive phenotype of *clc-3* mutants. Numbers at the base of each bar indicate the number of independent trials performed, with *n*=20 *C. elegans* per trial. Data are mean ± SEM. (D) Representative images showing the distribution of SNB-1 puncta at the cholinergic synapses along the DNC. Scale bar = 20 μm. Deletion of the PDZ bm prevents the rescue of the puncta-width defect observed in *clc-3* mutants, as indicated in the accompanying bar graphs. Numbers at the base of the bars indicate the number of *C. elegans* used for imaging. Data are mean ± SEM. Statistical analyses were performed using one-way ANOVA followed by Tukey’s post-hoc test.

### Mutants in *clc-3* and *nab-1* act in the same genetic pathway to regulate body bend amplitude

As the PDZ-binding motif has been implicated in CLC-3 function at the presynaptic cholinergic neurons, we sought to identify PDZ-domain containing interacting partners that could mediate downstream signalling. Given the well-established role of the actin cytoskeleton in regulating synaptic vesicle dynamics in both vertebrates and *C. elegans* (*62*), we focused our screen on proteins that both contain a PDZ domain and associate with the actin-cytoskeleton. Two candidates met these criteria; 1) ZOO-1, the sole member of the zona occludens family in *C. elegans* and 2) NAB-1(Neurabin-1), a known actin-binding protein. ZOO-1 has been shown to influence actin organization, potentially through interactions with other actin associated proteins, while NAB-1 modulates actin organisation via direct binding to F-actin (*43, 63, 64*).

We analysed *nab-1* and *zoo-1* mutants for defects in body bend amplitude and found that though *zoo-1* mutants displayed a modest increase in body-bend amplitude compared to WT *C. elegans*, the phenotype was significantly different from that of *clc-3* mutants (Figure 6A (left)). Mutants in *nab-1*, on the other hand, exhibited a defect that closely resembled the *clc-3* mutant phenotype (Figure 6A (right)). Based on this similarity, we chose *nab-1* mutants for further experiments and generated double mutants of *nab-1* and *clc-3* to test for potential genetic interaction between the two genes.

**Figure 6:**
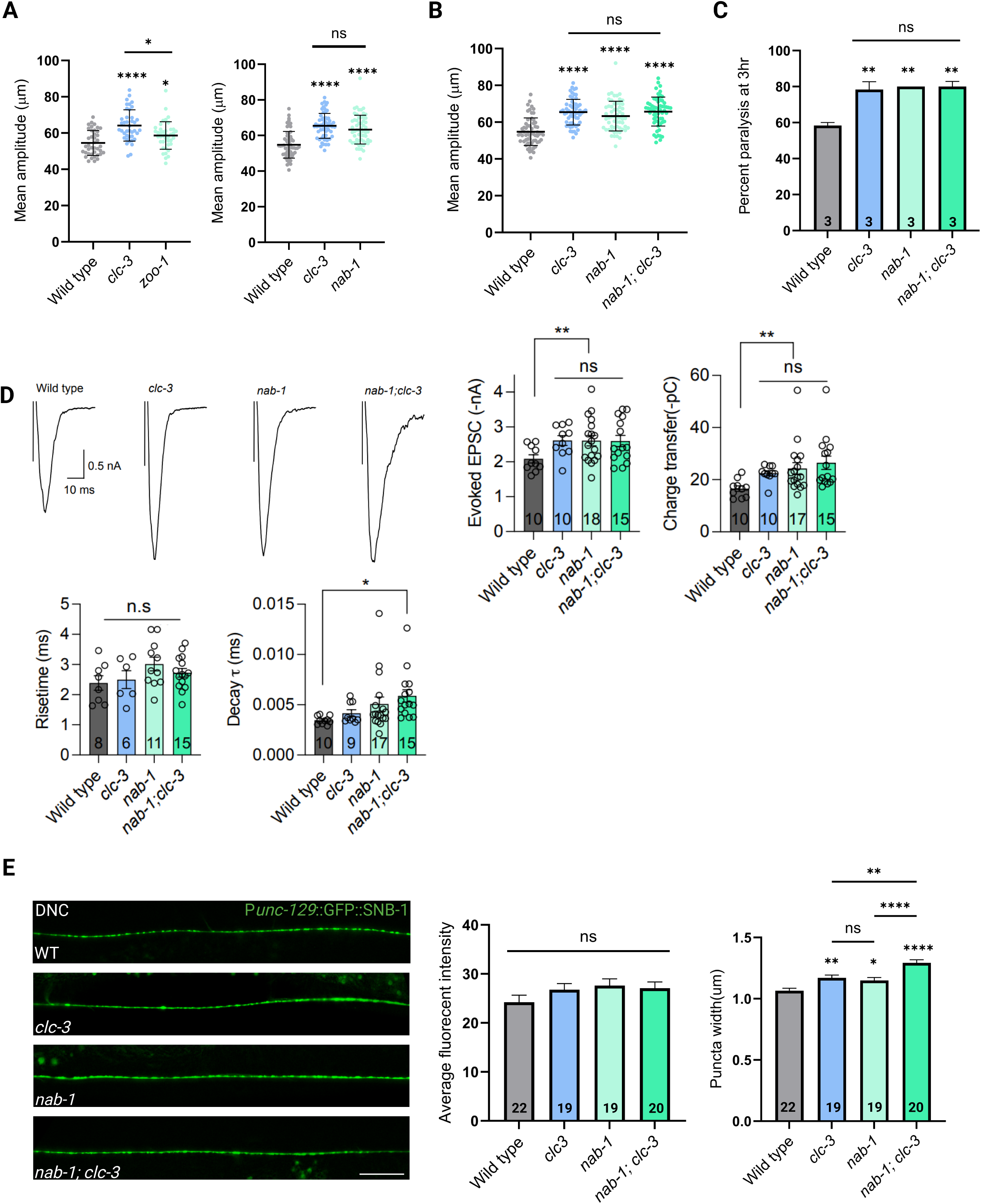
The genes *clc-3* and *nab-1* act in the same genetic pathway to regulate body bend amplitude (A) *nab-1* mutants exhibit increased body-bend amplitude, similar to *clc-3* mutants. Graphs show quantification of the mean body-bend amplitude in *zoo-1* (left) and *nab-1* mutants (right). *N*≥2; *n*=10. Data are mean ± SD. Statistical analyses were performed using one-way ANOVA followed by Tukey’s post-hoc test. (B) Graphs representing the quantification of mean body-bend amplitude show that *nab-1; clc-3* double mutants exhibit amplitudes comparable to those of the respective single mutants, suggesting that *nab-1* and *clc-3* genes may act in the same genetic pathway. *N*=3; *n*=10. Data are mean ± SD, \*\*\*\**p*<0.0001. Statistical analyses were performed using one-way ANOVA followed by Tukey’s post-hoc test. (C) Bar graph showing the percentage of *C. elegans* paralysed at the 180-minute time point following aldicarb exposure indicate that *nab-1; clc-3* double mutants display aldicarb hypersensitivity similar to that of either single mutant, further supporting a genetic interaction between *nab-1* and *clc-3*. Numbers at the base of each bar indicate the number of independent trials performed, with *n*=20 *C. elegans* per trial. Data are mean ± SEM. Statistical analyses were performed using one-way ANOVA followed by Tukey’s post-hoc test. (D) Representative traces of stimulus-evoked response recorded from the body wall muscles following electrical stimulation of the ventral cord. Bar graphs represent quantification of evoked current amplitude (left) and total charge transfer (right). *nab-1; clc-3* double mutants exhibit increases in both parameters comparable to those observed in respective single mutants. Numbers at the base of the bars indicate the number of *C. elegans* used for recordings. Data are mean ± SEM. (E) Representative images showing the distribution of SNB-1 puncta at the cholinergic synapses along the DNC (Scale bar = 20 μm). Bar graphs represent quantification of average fluorescent intensity (left) and puncta width (right). *nab-1* mutants exhibit a puncta width defect similar to *clc-3* mutants, while *nab-1; clc-3* double mutants show a further increase in puncta width compared to either single mutant. The average fluorescent intensity remains comparable across all genotypes. Numbers at the base of each bar indicate the total number of worms imaged. Data are mean ± SEM. Statistical analyses were performed using one-way ANOVA followed by Tukey’s post-hoc test.

First, we tested whether *nab-1* and *clc-3* function in the same genetic pathway to influence locomotion amplitude by quantifying the body-bend amplitude in *nab-1; clc-3* double mutants. We saw that the amplitude of the double mutants resembled that of both single mutants, indicating that the two genes likely act in the same genetic pathway (Figure 6B).

Next, to investigate whether *nab-1* single mutants and *nab-1; clc-3* double mutants exhibit synaptic transmission defects similar to those seen in *clc-3* mutants, we subjected them to aldicarb assays and electrophysiology studies. Both *nab-1* single mutants and *nab-1; clc-3* double mutants showed increased sensitivity to aldicarb, closely matching the hypersensitivity observed in *clc-3* mutants (Figures 6C and S6A). Additionally, electrophysiological recordings revealed a similar increase in the evoked postsynaptic currents across all three genotypes (Figure 6D). It should be noted that the decay in *nab-1; clc-3* double mutants is larger compared to the single mutants, indicating that NAB-1 may have an additional function in regulating slow release or synaptic vesicle replenishment. The spontaneous neurotransmitter release remained unaffected in all cases, suggesting a shared presynaptic defect across single and double mutants of *clc-3* and *nab-1* (Figures S6B and S6C). Consistent with the electrophysiology data, the width of SNB-1 puncta was increased in both *nab-1* and *nab-1; clc-3* mutants (Figure 6E). Notably, the *nab-1; clc-3* double mutants exhibited a further increase in puncta width compared to either single mutant, consistent with the prolonged decay in the evoked postsynaptic currents, suggesting that NAB-1 may have additional functions beyond its shared role with CLC-3 in regulating synaptic vesicle localization. The values of average fluorescent intensity and puncta density were similar across all genotypes (Figures 6E and S6D). Taken together, these findings support the idea that *nab-1* and *clc-3* function within the same genetic pathway to regulate the body-bend amplitude by mediating presynaptic organization.

### CLC-3 and NAB-1 interact with each other via the CLC-3 PDZ-binding motif

Imaging experiments revealed that NAB-1 and CLC-3 colocalise along the DNC (Figure 7A). To test whether the two proteins interact with each other physically to mediate normal body-bend amplitude, we performed Bimolecular Fluorescence Complementation (BiFC) assays using constructs where CLC-3 and NAB-1 were individually tagged with complementary YFP fragments and expressed under a cholinergic neuron-specific promoter (Illustrated in Figure 7B). A clear YFP signal was observed in the head cholinergic neurons and along the DNC when full-length CLC-3 and NAB-1 were co-expressed (Figures 7C and S7A). In contrast, no clear YFP signal was observed in *C. elegans* expressing CLC-3 lacking the PDZ-binding motif along with NAB-1, indicating that the interaction of the two proteins is dependent on this motif (Figures 7C and S7A).

**Figure 7:**
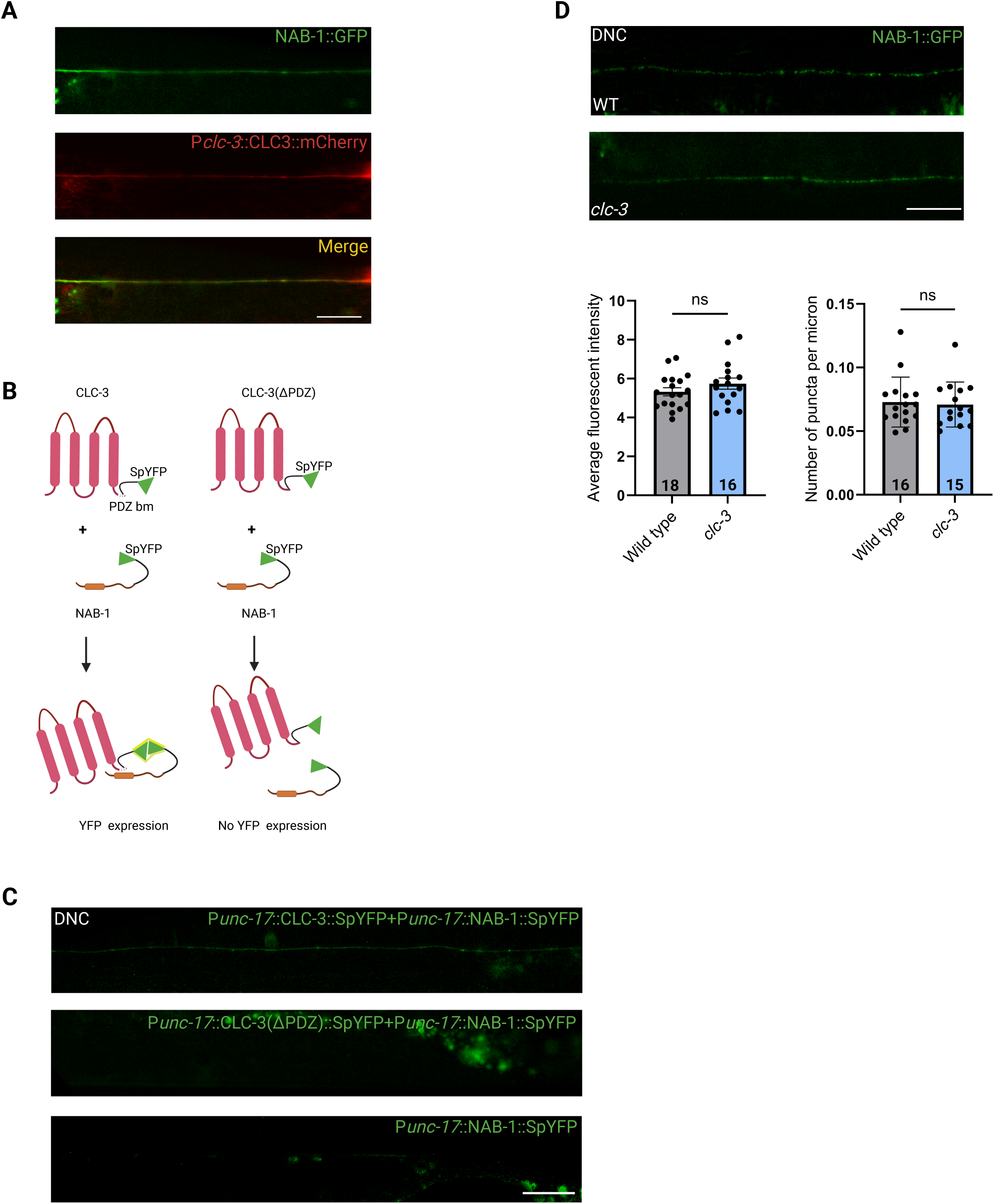
CLC-3 and NAB-1 physically interact with each other via the CLC-3 PDZ-binding motif (A) Representative images showing NAB-1::GFP and CLC-3::mCherry expression along the DNC. CLC-3 expression overlaps with NAB-1 along the DNC. Scale bar = 20 μm. (B) Schematic illustrating the possible outcomes of the BiFC assay. CLC-3 (pink) is fused to the C-terminal half of YFP, while the C-terminus of NAB-1(orange) is fused to the N-terminal half of YFP. Both YFP fragments (green) are linked to their respective proteins via flexible linkers (black). The PDZ-binding motif of CLC-3 is represented by white circles at its C-terminus. (C) Representative images of the DNC of WT animals expressing either P*unc-17*::CLC-3::SpYFP and P*unc-17*::NAB-1::SpYFP together, P*unc-17*::CLC-3(ΔPDZ)::SpYFP and P*unc-17*::NAB-1::SpYFP together, or only P*unc-17*::NAB-1::SpYFP. The interaction between NAB-1 and full-length CLC-3 leads to reconstitution of YFP fluorescence along the DNC in cholinergic neurons, but no fluorescence is detected in the absence of the PDZ bm or when P*unc-17*::NAB-1::SpYFP is expressed alone. Scale bar = 20 μm, *n*=10-17. (D) Representative images showing NAB-1::GFP expression along the DNC in WT and *clc-3* mutant *C. elegans.* Scale bar = 20 μm. Bar graphs showing quantification of average fluorescent intensity (left) and puncta density (right) indicate that NAB-1 expression in mutants is similar to that of WT *C. elegans*. Numbers at the base of each bar indicate the total number of worms imaged. Data are mean ± SEM. Statistical analyses were performed using unpaired Student’s *t* test.

To determine whether CLC-3 influences NAB-1 expression, we visualised NAB-1::GFP in the *clc-3* mutant background. Imaging analyses revealed that the expression and localization of NAB-1 in *clc-3* mutants were similar that seen in WT animals (Figure 7D).

Together, these findings indicate that CLC-3 and NAB-1 interact with each other in cholinergic neurons via the CLC-3 PDZ-binding motif and that this interaction likely underlies their shared role in modulating body-bend amplitude and synaptic signaling.

## Discussion

Neurons across vertebrates and invertebrates express a rich repertoire of CAMs, which are essential for diverse aspects of nervous system function (*1, 65–69*). Mutations in CAMs can result in a spectrum of neuronal and synaptic defects, ranging from gross structural abnormalities, such as severe disruptions in axon development and guidance, to more subtle yet functionally significant impairments in synaptic organisation and plasticity (*70–73*).

Claudins, best known for their role in tight junctions, represent a class of CAMs whose expression and function within the neurons and at the synapse remain relatively underexplored (*42*). Notably, claudins exhibit several structural and functional parallels with classical neuronal CAMs; 1) Like other synaptic and neuronal CAMs, claudins possess extracellular and intracellular domains that enable interactions with extracellular partners, including other claudins and intracellular signalling molecules. These interactions allow claudins to participate in signalling cascades that regulate critical cellular processes such cell morphology and polarity (*21, 61, 74–76*). 2) Similar to other neuronal CAMs, claudins are components of complexes that anchor them to the cytoskeleton, primarily through interactions with actin-binding proteins mediated by their C-terminal PDZ-binding motif (*77–80*). 3) Beyond the tight junctions, claudins have also been shown to interact with non-tight junction proteins, including other

CAMs and gap junction proteins. For example, Claudin-7 has been shown to transiently interact with Connexin 43 and E-cadherin in murine mammary glands during post-natal development (*80*). Similar types of interactions have been reported for other CAMs. For instance, the *Drososphila* E-cadherin has been shown to directly interact with the gap junction protein Innexin2 (*81*). Although these interactions have been reported in non-neuronal tissues, they raise the possibility that claudins, like other NCAMs, may participate in diverse scaffolding and regulatory complexes in the nervous system. 4) Lastly, from a pathological standpoint, defects in both claudins and known neuronal CAMs have been linked with several nervous system disorders. For example, Claudin-5 dysfunction has been associated with disorders such as Alzheimer’s, multiple sclerosis and schizophrenia (*82–85*). Likewise, changes in the expression of certain neuronal CAMs, such as NCAMs, have been associated with disorders such as Alzheimer’s, schizophrenia and bipolar disorder (*70, 86–89*).

Despite these important insights, the study of claudin function has historically been biased towards epithelial and endothelial contexts. In contrast, their expression and function within neurons have remained largely overlooked. This neglect may, in part, stem from the absence of classical tight junctions in neurons as well as the possibility that claudin expression in the vertebrate nervous system is more restricted, either confined to specific brain regions or neuronal subtypes, or expressed at levels that are below the detection threshold of conventional screening approaches. Invertebrate model systems, particularly *C. elegans,* offer several advantages that circumvent many of the challenges associated with vertebrate models, including a simple nervous system composed of a finite number of neurons with a well-mapped neural connectome, and the ease of genetic manipulation (*90, 91*). In accordance with this, and perhaps unsurprisingly, few claudin-like molecules in *C. elegans*, namely HIC-1, HPO-30 and NSY-4, have been shown to have neuronal and synaptic roles (*43–45, 47*). These findings lend further support to the hypothesis that claudins may play important roles in neurons.

Leveraging this simple yet powerful model system, we systematically examined the neuronal expression of 17 claudin-like proteins in *C. elegans*. We found that nine of these proteins were expressed in neurons, among which a candidate called *clc-3* stood out for its robust expression that was restricted to the neurons. Functional analyses revealed that *clc-3* is involved in regulating locomotion in *C. elegans*, potentially through a presynaptic mechanism. Mutants in *clc-3* exhibit an increased sinusoidal wave amplitude with no accompanying changes in overall movement speed. This phenotype was rescued by reintroducing *clc-3* under its endogenous promoter or by specifically expressing it in cholinergic neurons in the mutant background, implicating a neuron-specific role in regulating locomotor output. The sinusoidal wave motion in *C. elegans* is known to be regulated by the co-ordinated excitation and inhibition of body wall muscles by ventral cord motor neurons. A disruption in this excitation-inhibition balance can alter the movement of worms. Hypersensitivity of *clc-3* mutants towards the chemical aldicarb suggested that similar alterations in excitation-inhibition balance may underlie the amplitude defect in *clc-3* mutants.

Electrophysiological recordings from the body wall muscles of *clc-3* mutants revealed a specific presynaptic role of the molecule. While both the frequency and amplitude of spontaneous excitatory and inhibitory postsynaptic currents, as well as responses to exogenous ACh remained unaltered in *clc-3* mutants, the evoked post-synaptic currents were significantly higher. In other words, electrical stimulation of the ventral cord caused an increase in the postsynaptic currents at the NMJ in *clc-3* mutants.

These findings suggest that while the machinery supporting spontaneous neurotransmission and postsynaptic receptor sensitivity is intact in *clc-3* mutants, activity-dependent neurotransmitter release is enhanced, pointing towards a presynaptic defect, possibly in the neurotransmitter vesicle release probability.

Imaging analyses of fluorescently tagged markers in *clc-3* mutants provided anatomical validation for electrophysiological findings. We observed that the active zone morphology at cholinergic synapses, marked by P*unc-129*::SYD-2::YFP and postsynaptic ACR-16 receptor expression appeared normal in *clc-3* mutants. Additionally, the neuromuscular junction, visualised using P*him-4*::MB::YFP, was structurally intact and showed no signs of developmental defects. To rule out the possibility that the deeper body-bend amplitude stemmed from changes in muscle tone, we examined MYO-3::GFP expression, which also appeared unaffected. Together, these results indicate that the synaptic architecture in *clc-3* mutants is grossly normal, consistent with the unaltered spontaneous post-synaptic currents and responses to exogenous ACh. However, analyses of the synaptic vesicle marker SNB-1 revealed abnormal aggregation of SNB-1 puncta in *clc-3* mutants. Specifically, there was an increase in the punctal width, without accompanying changes in puncta density, indicating a potential defect in the organization of the vesicle pool (Illustrated in Figure 8). These alterations correlated with the increased evoked postsynaptic currents, suggesting that the disrupted vesicle clustering may lead to an enhanced availability of synaptic vesicles during activity-dependent release, thereby enhancing neurotransmitter output in the presence of an stimulus.

**Figure 8:**
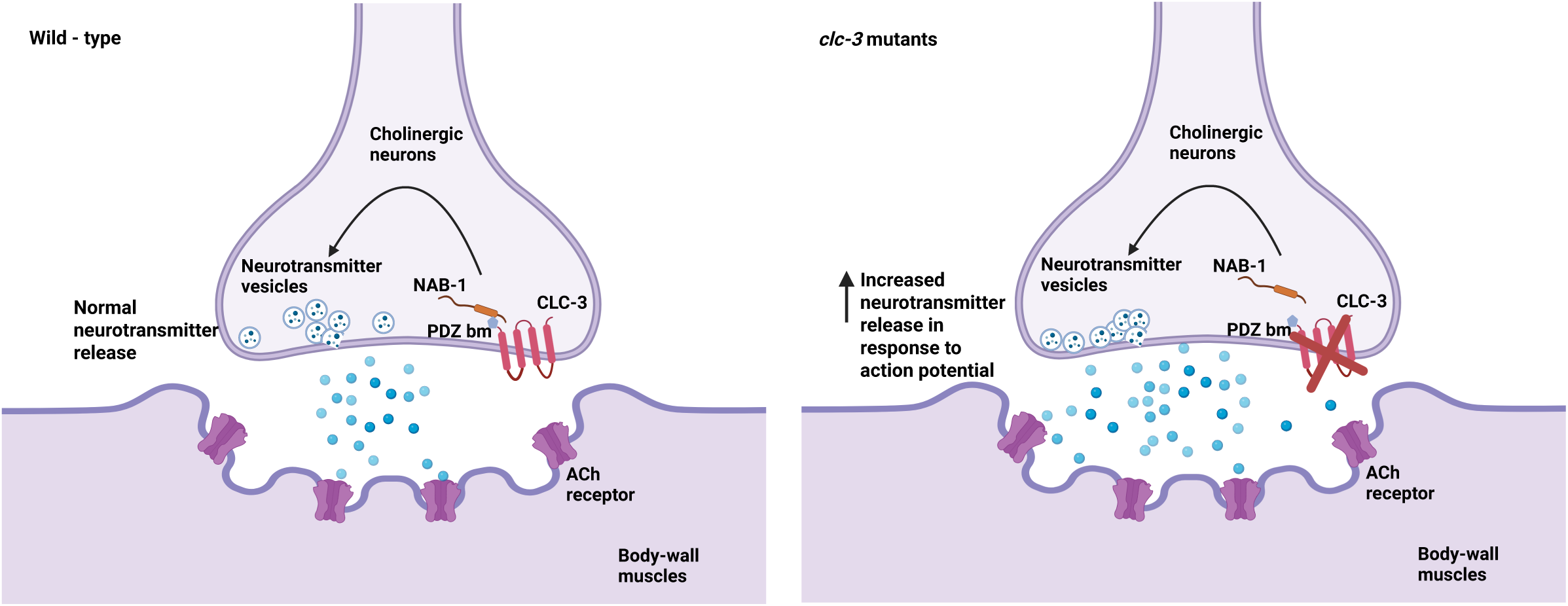
Illustrative model depicting the role of CLC-3 in cholinergic neurons The schematic summarises the proposed mechanism of CLC-3 function in cholinergic neurons. Our data indicate that CLC-3 is expressed at the presynaptic membrane in cholinergic neurons, where it interacts with the actin-binding protein NAB-1 via its C-terminal PDZ-binding motif and regulates the organization of the synaptic vesicle pool. In WT *C. elegans*, CLC-3 functions to negatively regulate the neurotransmitter release at the neuromuscular synapses, likely by maintaining proper vesicle clustering at the presynapse (left). Loss of CLC-3 results in increased evoked postsynaptic currents, possibly due to dysregulated vesicle release, which may contribute to the body-bend amplitude defect observed in mutants (right).

In support of a presynaptic role for CLC-3, we identified a direct physical interaction between CLC-3 and an actin-binding protein, NAB-1. This interaction occurs via the PDZ-binding motif of CLC-3 and seems functionally relevant. *nab-1; clc-3* double mutants display phenotypes comparable to those observed in either single mutant, including increased body bend amplitude, aldicarb hypersensitivity, elevated evoked release and increased SNB1 puncta width, suggesting that CLC-3 and NAB-1 act in a common genetic pathway.

Our findings point towards a presynaptic defect in fine-tuning activity-dependent neurotransmitter release in *clc-3* mutants, without affecting basal synaptic transmission or postsynaptic receptor function. This role of CLC-3 in regulating synaptic organization is reminiscent of other neuronal CAMs that fine-tune synaptic strength and architecture in both *C. elegans* and vertebrates. For example, mutants of SYG-1, a CAM belonging to the immunoglobulin family in *C. elegans*, exhibit defects in synaptic vesicle clustering, where vesicles cluster ectopically and fail to accumulate at synaptic sites in HSNL neuron (*10*). Intriguingly, these mutants also show increased body bend amplitude during locomotion, as reported in another study (*92*). In vertebrates, the proteolytic cleavage of Neuroligin-1, a post-synaptic CAM, reduces the probability of neurotransmitter release at the presynapse by destabilising its presynaptic partner, neurexin-1b, highlighting how structural modifications of CAMs can serve as a mechanism for influencing synaptic transmission (*93*). In this context, a compelling feature that could make claudins such as CLC-3 well suitable for fine-tuning synaptic function is their capacity to undergo diverse post-translational modifications (PTMs), such as phosphorylation, palmitoylation, and glycosylation, particularly within their C-terminus. These modifications can impact claudin localization, stability, and interaction with binding partners. Because PTMs are rapid and reversible, they allow claudins to be dynamically regulated in response to changes in the external environment, a key requirement for modulating synaptic strength in an activity-dependent manner (*22, 94, 95*).

Taken together, our findings provide further evidence for the expanded role of claudin-like molecules in *C. elegans* beyond their canonical barrier functions. By showing that CLC-3 acts as a modulator of presynaptic organization and motor circuit output, we highlight how claudins in *C. elegans* can integrate changes in synaptic signalling with whole-animal behaviour. Importantly, these findings underscore how subtle synaptic alterations, often overlooked in favour of gross anatomical defects, can offer meaningful insights into the fine-tuning of evolutionarily conserved behavioural outputs.

## Acknowledgements

We thank Josh Kaplan for kindly providing marker strains. Some strains were provided by CGC, which is funded by NIH Office of Research Infrastructure Programs (P40 OD010440). All illustrations were created in BioRender. We are grateful to Professors Deepak Nair and Arnab Barik for their valuable feedback during annual project presentations, and to Professor Abhishek Bhattacharya for his insightful comments during joint lab meetings. We thank Srushti Bhasme and Nijin J for assistance with experiments. We also thank Palagiri Suresh for routine help and the members of Kavita Babu’s lab for suggestions and critique on the manuscript.

## Author Contributions

AA: design and execution of experiments, data analyses, manuscript writing/editing and supervision, AR: execution of experiments and data analyses, HL: execution of experiments and data analyses, AS: execution of experiments, JJ: execution of experiments, HK: execution of experiments, ZH: supervision and manuscript editing, and KB: initial idea, supervision, funding acquisition, and manuscript writing/editing.

