## Supplemental Information for "The claudin-like molecule CLC-3 regulates neuromuscular function in *Caenorhabditis elegans* by modulating cholinergic signalling"

**Supplementary information**

**Table S1 (List of strains)**

| Strain name | Strain description | CGC strain | Outcross status | Figure number |
|---|---|---|---|---|
| BAB7001 | IndEx7001 [P*clc-1*::GFP] | This study | | 1 |
| BAB7002 | IndEx7002 [P*clc-2*::GFP] | This study | | 1 |
| BAB7003 | IndEx7003 [P*clc-3*::GFP] | This study | | 1 |
| BAB7004 | IndEx7004 [P*clc-4*::GFP] | This study | | 1 |
| BAB7005 | IndEx7005 [P*clc-5*::GFP] | This study | | 1 |
| BAB7006 | IndEx7006 [P*clc-6*::GFP] | This study | | 1 |
| BAB7007 | IndEx7007 [P*clc-7*::GFP] | This study | | 1 |
| BAB7008 | IndEx7008 [P*clc-8*::GFP] | This study | | 1 |
| BAB7009 | IndEx7009 [P*clc-9*::GFP] | This study | | 1 |
| BAB7010 | IndEx7010 [P*clc-11*::GFP] | This study | | 1 |
| BAB7011 | IndEx7011 [P*clc-14*::GFP] | This study | | 1 |
| BAB7012 | IndEx7012 [P*clc-32*::GFP] | This study | | 1 |
| BAB7013 | IndEx7013 [P*nsy-4*::GFP] | This study | | 1 |
| BAB7014 | IndEx7014 [P*vab-9*::GFP] | This study | | 1 |
| BAB7015 | IndEx7015 [P*hpo-30*::GFP] | This study | | 1 |
| BAB7016 | IndEx7016 [P*hic-1*::GFP] | This study | | 1 |
| BAB7017 | IndEx7017 [P*F53B3.5*::GFP] | This study | | 1 |
| BAB7018 | *clc-3 (ok1584)* | RB1394 | 4X | 2-6 |
| BAB7019 | IndEx7018 [P*clc-3*::CLC-3::T2A::GFP (I); *clc-3*] | This study | | 2, 5 |
| BAB7020 | IndEx7019 [P*clc-3*::CLC-3::T2A::GFP (II); *clc-3*] | | | |
| BAB7021 | IndEx7020 [P*unc-17*::mCherry; P*clc-3*::GFP] | This study | | 2 |
| BAB7022 | IndEx7021 [P*unc-25*::mCherry; P*clc-3*::GFP] | | | |
| BAB7023 | IndEx7022 [P*unc-17*::CLC-3::T2A::GFP; *clc-3*] | This study | | 2, 3, 5 |
| BAB7024 | IndEx7023 [P*unc-25*::CLC-3::T2A::GFP; *clc-3*] | This study | | 2 |
| BAB7025 | IndEx7024 [P*let-413*::CLC-3::T2A::GFP; *clc-3*] | This study | | 2 |
| BAB7026 | IndEx7025 [P*myo-3*::CLC-3::T2A::GFP; *clc-3*] | This study | | 2 |
| BAB7027 | P*unc-129*::GFP::SNB-1 | KP3814 (*nuIs152*) | 3X | 4, 5, 6 |

| | | | | |
|---|---|---|---|---|
| BAB7028 | P*unc-129*::GFP::SNB-1; *clc-3* | This study | | 4, 5, 6 |
| BAB7029 | IndEx7026 [P*clc-3*::CLC-3::mCherry; BAB7027] | This study | | 4 |
| BAB7030 | IndEx7027 [P*unc-17*::CLC-3::T2A::mCherry; BAB7028] | This study | | 4, 5, 6 |
| BAB7031 | P*unc-129*::SYD-2::YFP | *nuIs159* | 3X | 4 |
| BAB7032 | P*unc-129*::SYD-2::YFP; *clc-3* | This study | | 4 |
| BAB7031 | P*myo-3*::ACR-16::GFP | *nuIs299* | 2X | 4 |
| BAB7033 | P*myo-3*::ACR-16::GFP; *clc-3* | This study | | 4 |
| BAB7034 | IndEx7028 [Punc-17::mCherry] | This study | | 4 |
| BAB7035 | IndEx7029 [P*unc-17*::mCherry; *clc-3*] | This study | | 4 |
| BAB7036 | P*him-4*::MB::YFP | RP247 | | 4 |
| BAB7037 | P*him-4*::MB::YFP; *clc-3* | This study | | 4 |
| BAB7038 | P*myo-3*::GFP::MYO-3 | RW1596 | 3X | 4 |
| BAB7039 | P*myo-3*::GFP::MYO-3; *clc-3* | This study | | 4 |
| BAB7040 | IndEx7030 [P*clc-3*::CLC-3(ΔPDZ)::T2A::GFP; *clc-3*] | This study | | 5 |
| BAB7041 | IndEx7031 [P*clc-3*::CLC-3(ΔPDZ)::T2A::mCherry; BA7028] | This study | | 5 |
| BAB7042 | IndEx7032 [P*unc-17*::CLC-3(ΔPDZ)::T2A::mCherry; BAB7028] | This study | | 5 |
| BAB7043 | *zoo-1 (gk404)* | VC896 | 4X | 6 |
| BAB7044 | *nab-1 (ok943)* | RB1017 | 4X | 6 |
| BAB7045 | *nab-1; clc-3* | This study | | 6 |
| BAB7046 | *nab-1*; BAB7027 | This study | | 6 |
| BAB7047 | *nab-1; clc-3;* BAB7027 | This study | | 6 |
| BAB7048 | NAB-1::GFP | ZM1385 (*hpIs66*) | 3X | 7 |
| BAB7049 | IndEx7033 [P*clc-3*::CLC-3::mCherry; BAB7048] | This study | | 7 |
| BAB7050 | NAB-1::GFP; *clc-3* | This study | | 7 |
| BAB7051 | IndEx7034 [P*unc-17*::CLC-3::SpYFP + P*unc-17*::NAB-1::SpYFP] | This study | | 7 |
| BAB7052 | IndEx7035 [P*unc-17*::CLC-3(ΔPDZ)::SpYFP + P*unc-17*::NAB-1::SpYFP] | This study | | 7 |

| | | | | |
|---|---|---|---|---|
| BAB7053 | IndEx7036 [P*unc-17*::CLC-3::SpYFP] | This study | | 7 |
| BAB7054 | IndEx7037 [P*unc-17*::NAB-1::SpYFP] | This study | | 7 |

**Table S2 (List of genotyping primers)**

| **Strain** | **Primer** | **Description** |
|---|---|---|
| *clc-3 (ok1584)* | AR015-TCTGCGTGTTCTGGTGGTTA<br><br>AR016-TGGGGGATGTCTTCTTAACG | External Forward<br><br>External Reverse |
| *nab-1 (ok943)* | AS019-ATCGCAATTTTCTCCATTCG<br><br>AS020-GCGTACTTCTTTCGGAGTGG | External Forward<br><br>External Reverse |

**Table S3 (List of plasmids)**

| **Plasmid** | **Construct** | **Backbone** |
|---|---|---|
| pBAB7001 | P*clc-1*::GFP | pPD95.75 |
| pBAB7002 | P*clc-2*::GFP | pPD95.75 |
| pBAB7003 | P*clc-3*::GFP | pPD95.75 |
| pBAB7004 | P*clc-4*::GFP | pPD95.75 |
| pBAB7005 | P*clc-5*::GFP | pPD95.75 |
| pBAB7006 | P*clc-6*::GFP | pPD95.75 |
| pBAB7007 | P*clc-7*::GFP | pPD95.75 |
| pBAB7008 | P*clc-8*::GFP | pPD95.75 |
| pBAB7009 | P*clc-9*::GFP | pPD95.75 |
| pBAB7010 | P*clc-11*::GFP | pPD95.75 |
| pBAB7011 | P*clc-14*::GFP | pPD95.75 |
| pBAB7012 | P*clc-32*::GFP | pPD95.75 |
| pBAB7013 | P*nsy-4*::GFP | pPD95.75 |
| pBAB7014 | P*vab-9*::GFP | pPD95.75 |
| pBAB7015 | P*hpo-30*::GFP | pPD95.75 |
| pBAB7016 | P*hic-1*::GFP | pPD95.75 |
| pBAB7017 | P*F53B3.5*::GFP | pPD95.75 |
| pBAB7018 | P*clc-3*::CLC-3::T2A::GFP | pPD95.75 |
| pBAB7019 | P*unc-17*::mCherry | pPD95.75 |
| pBAB7020 | P*unc-17*::CLC-3::T2A::GFP | pPD95.75 |
| pBAB7021 | P*unc-25*::CLC-3::T2A::GFP | pPD95.75 |
| pBAB7022 | P*let-413*::CLC-3::T2A::GFP | pPD95.75 |
| pBAB7023 | P*myo-3*::CLC-3::T2A::GFP | pPD95.75 |
| pBAB7024 | P*clc-3*::CLC-3::mCherry | pPD95.75 |
| pBAB7025 | P*unc-17*::CLC-3::T2A::mCherry | pPD95.75 |
| pBAB7026 | P*clc-3*::CLC-3(ΔPDZ)::T2A::GFP | pPD95.75 |
| pBAB7027 | P*clc-3*::CLC-3(ΔPDZ)::T2A::mCherry | pPD95.75 |
| pBAB7028 | P*unc-17*::CLC-3(ΔPDZ)::T2A::mCherry | pPD95.75 |
| pBAB7029 | P*unc-17*::CLC-3::SpYFP(VC155) | pPD95.75 |
| pBAB7030 | P*unc-17*::CLC-3(ΔPDZ)::SpYFP(VC155) | pPD95.75 |
| pBAB7031 | P*unc-17*::NAB-1::SpYFP(VN173) | pPD95.75 |
| pBAB7032 | P*unc-17*::NAB-1::T2A::mCherry | pPD95.75 |

**Table S4 (List of primers used for cloning)**

| Plasmid | Primers | Description |
|---|---|---|
| P*clc-1*::GFP | AA047-ATGGATACGCTAACAACTTGGAAATGAAATAAGCTTGCATCGCTCTGCATTGATTTGATG | Forward primer for amplifying *clc-1* promoter from genomic DNA |
| | AA060-TCCTTTGGCCAATCCCGGGGATCCTCTAGAGTCGACCTGCATTGTGAACCTGAAACATATGGT | Reverse primer for amplifying *clc-1* promoter from genomic DNA |
| P*clc-2*::GFP | AA076-GAAATGAAATAAGCTTGCATGCAAGAGAATTATACAGTTGATATATATGATTTTCAAG | Forward primer for amplifying *clc-2* promoter from genomic DNA |
| | AA085-CTCATTTTTTCTACCGGTACCTTCTGCAAAGAAGACGTAAATTAATACAAG | Reverse primer for amplifying *clc-2* promoter from genomic DNA |
| P*clc-3*::GFP | AA049-ATGGATACGCTAACAACTTGGAAATGAAATAAGCTTGCATAAGTGGCCGGATGAGATGTA | Forward primer for amplifying *clc-3* promoter from genomic DNA |
| | AA062-TCCTTTGGCCAATCCCGGGGATCCTCTAGAGTCGACCTGCCATTTTGAATTCCTCGTAATCAGAC | Reverse primer for amplifying *clc-3* promoter from genomic DNA |
| P*clc-4*::GFP | AA050-ATGGATACGCTAACAACTTGGAAATGAAATAAGCTTGCATTCAAGTTTGGACAAGCTATAGAGAAA | Forward primer for amplifying *clc-4* promoter from genomic DNA |

| | | |
|---|---|---|
| | AA063-TCCTTTGGCCAATCCCGGGGATCCTCTAGAGTCGACCTGCCATCCAGGATCACTGAAACATAA | Reverse primer for amplifying *clc-4* promoter from genomic DNA |
| P*clc-5*::GFP | AA077-ATGGATACGCTAACAACTTGGAAATGAAATAAGCTTGCATTTGTTCTGAGCATGTTTGATTTTT | Forward primer for amplifying *clc-5* promoter from genomic DNA |
| | AA086-CCAATCCCGGGGATCCTCTAGAGTCGACCTGC TCCCAATGCCATTCACACTA | Reverse primer for amplifying *clc-5* promoter from genomic DNA |
| P*clc-6*::GFP | AA052-ATGGATACGCTAACAACTTGGAAATGAAATAAGCTTGCATAGCCGCTTCGTAATACCTCA | Forward primer for amplifying *clc-6* promoter from genomic DNA |
| | AA065-TCCTTTGGCCAATCCCGGGGATCCTCTAGAGTCGACCTGCTCTGCAAACGAATAAAAACATGA | Reverse primer for amplifying *clc-6* promoter from genomic DNA |
| P*clc-7*::GFP | AA082-ATGGATACGCTAACAACTTGGAAATGAAATAAGCTTGCATTTGTCCTCATTGTTGCTTCG | Forward primer for amplifying *clc-7* promoter from genomic DNA |
| | AA091-TCCTTTGGCCAATCCCGGGGATCCTCTAGAGTCGACCTGC GGTTGCGTCTGAAACATATTCGA | Reverse primer for amplifying *clc-7* promoter from genomic DNA |
| P*clc-8*::GFP | AA081-GAAATGAAATAAGCTTGCATGCATCTATTGGAAATTCAAATATTTACGGATGG | Forward primer for amplifying *clc-8* promoter from genomic DNA |

| | | |
|---|---|---|
| | AA090-CTCATTTTTTCTACCGGTACCCCCATTGGCTCTGAAAATGTGAAATTTTC | Reverse primer for amplifying *clc-8* promoter from genomic DNA |
| P*clc-9*::GFP | AA054-ATGGATACGCTAACAACTTGGAAATGAAATAAGCTTGCATTAGAAAATTGTGTTTAAGCTCTTCCA | Forward primer for amplifying *clc-9* promoter from genomic DNA |
| | AA067-TCCTTTGGCCAATCCCGGGGATCCTCTAGAGTCGACCTGCCTGAAAAATTAAATTACAGCAGGATA | Reverse primer for amplifying *clc-9* promoter from genomic DNA |
| P*clc-11*::GFP | AA080-ATGGATACGCTAACAACTTGGAAATGAAATAAGCTTGCATTGGAGCAGCACAAAACCATA | Forward primer for amplifying *clc-11* promoter from genomic DNA |
| | AA089-CCAATCCCGGGGATCCTCTAGAGTCGACCTGCTCATTCTCGGTGAGTGGAAA | Reverse primer for amplifying *clc-11* promoter from genomic DNA |
| P*clc-14*::GFP | AA078-ATGGATACGCTAACAACTTGGAAATGAAATAAGCTTGCATCCAGTCACATGTCCTCCAGA | Forward primer for amplifying *clc-14* promoter from genomic DNA |
| | AA087-TCCTTTGGCCAATCCCGGGGATCCTCTAGAGTCGACCTGC CGACAAACATTATTCCAAAACCTTAG | Reverse primer for amplifying *clc-14* promoter from genomic DNA |
| P*clc-32*::GFP | AA079-ATGGATACGCTAACAACTTGGAAATGAAATAAGCTTGCATGCCGTCAGATATTTTCAACGA | Forward primer for amplifying *clc-32* promoter from genomic DNA |

| | | |
|---|---|---|
| | AA088-CAATCCCGGGGATCCTCTAGAGTCGACCTGCTGACAGCCGACATTTCAAGT | Reverse primer for amplifying *clc-32* promoter from genomic DNA |
| P*nsy-4*::GFP | AA043-ATGGATACGCTAACAACTTGGAAATGAAATAAGCTTGCATATCTGCCAGCAGGTTTTCAT | Forward primer for amplifying *nsy-4* promoter from genomic DNA |
| | AA056-TCCTTTGGCCAATCCCGGGGATCCTCTAGAGTCGACCTGCTCGACGCTGAAATTTTTAAGG | Reverse primer for amplifying *nsy-4* promoter from genomic DNA |
| P*hpo-30*::GFP | AA075-ATGGATACGCTAACAACTTGGAAATGAAATAAGCTTGCATAACGAACAATTTGCGGAGAG | Forward primer for amplifying *hpo-30* promoter from genomic DNA |
| | AA084-CCAATCCCGGGGATCCTCTAGAGTCGACCTGCATGGCATTCCGTGGTTACTC | Reverse primer for amplifying *hpo-30* promoter from genomic DNA |
| P*hic-1*::GFP | AA045-ATGGATACGCTAACAACTTGGAAATGAAATAAGCTTGCATTTTTTCCTGCGGAGATATGG | Forward primer for amplifying *hic-1* promoter from genomic DNA |
| | AA058-TCCTTTGGCCAATCCCGGGGATCCTCTAGAGTCGACCTGCCCGGAAAAATACTTACAGAAAAGA | Reverse primer for amplifying *hic-1* promoter from genomic DNA |
| P*F53B3.5*::GFP | AA083-ATGGATACGCTAACAACTTGGAAATGAAATAAGCTTGCATCACAACGGTGTTCTACTCCAAA | Forward primer for amplifying *F53B3.5* promoter from genomic DNA |

| | | |
|---|---|---|
| | AA092-AATCCCGGGGATCCTCTAGAGTCGACCTGCTGACACGGTGAGGAAAACATTG | Reverse primer for amplifying *F53B3.5* promoter from genomic DNA |
| P*clc-3*::CLC-3::T2A::GFP | AA271-TTACGAGGAATTCAAAATGGCTATGCAATTCACCGGCCTACAGGTTTG | Forward primer for cloning CLC-3 genomic DNA along with T2A sequence upstream of GFP in pBAB7003 |
| | AA272-CGACGTCACCGCATGTTAGCAGACTTCCTCTGCCCTCTACCGGTACGTAAACTTCAGTTACTGGCGGG | Reverse primer for cloning CLC-3 genomic DNA along with T2A sequence upstream of GFP in pBAB7003 (overhang contains one half of T2A sequence) |
| | AA273-CTAACATGCGGTGACGTCGAGGAGAATCCTGGCCCAGAAAAAATGAGTAAAGGAGAAGAACTTTTC | Forward primer for amplifying backbone of vector pBAB7003 (overhang contains second half of T2A sequence) |
| | AA274-TTTGAATTCCTCGTAATCAGACGAACTT | Reverse primer for amplifying backbone of vector pBAB7003 |
| P*unc-17*::mCherry | AA275-GAAATGAAATAAGCTTGCATGCGCACACACACACATCCTCTCTTC | Forward primer for cloning *unc-17* promoter upstream to mCherry in PpD95.75 backbone |
| | AA276-CCTTTGAGACCATACCGGTACCCTTGTTTTAGTAGGTTACTATTTTGAACAAG | Reverse primer for cloning *unc-17* promoter upstream to mCherry in PpD95.75 backbone |
| P*unc-17*::CLC-3::T2A::GFP | AA277-GAAATGAAATAAGCTTGCATGCGCACACACACACATCCTCTCTTC | Forward primer for replacing *clc-3* promoter with *unc-17* promoter in pBAB7018 |

| | | |
|---|---|---|
| | AA278-GAATTGCATAGCCATGGTACCCTTGTTTTAGTAGGTTACTATTTTGAACAAG | Reverse primer for replacing *clc-3* promoter with *unc-17* promoter in pBAB7018 |
| P*unc-25*::CLC-3::T2A::GFP | AA279-GAAATGAAATAAGCTTGCATGCAGGATTTGATAGATGAGGAAAATCAAAG | Forward primer for replacing *clc-3* promoter with *unc-25* promoter in pBAB7018 |
| | AA280-GAATTGCATAGCCATGGTACCTTTTGGCGGTGAACTGAGCTTT<br>TC | Reverse primer for replacing *clc-3* promoter with *unc-25* promoter in pBAB7018 |
| P*let-413*::CLC-3::T2A::GFP | AA281-GAAATGAAATAAGCTTGCATGCTTGATTCTAGTTTACTTTAAAAATGAATATACCG | Forward primer for replacing *clc-3* promoter with *let-413* promoter in pBAB7018 |
| | AA282-GAATTGCATAGCCATGGTACCTGTGGCGATATATGAAGTTGATCG | Reverse primer for replacing *clc-3* promoter with *let-413* promoter in pBAB7018 |
| P*myo-3*::CLC-3::T2A::GFP | AA283-GAAATGAAATAAGCTTGCATGCGGTCTGGCCGCAAAAAGGTTTATG | Forward primer for replacing *clc-3* promoter with *myo-3* promoter in pBAB7018 |
| | AA284-GAATTGCATAGCCATGGTACCTTCTAGATGGATCTAGTGGTCGTG | Reverse primer for replacing *clc-3* promoter with *myo-413* promoter in pBAB7018 |
| P*clc-3*::CLC-3::mCherry | AA285-GAAATGAAATAAGCTTGCATGCGCACGTATAACCACTGGTAATTAG | Forward primer for cloning *clc-3* promoter and genomic DNA upstream to mCherry in PpD95.75 backbone |

| | | |
|---|---|---|
| | AA286-CCTTTGAGACCATACCGGTACCGTAAACTTCAGTTACTGGCGGG | Reverse primer for cloning *clc-3* promoter and genomic DNA upstream to mCherry in PpD95.75 backbone |
| P*unc-17*::CLC-3::T2A::mCherry | AA208-AGAATCCTGGCCCAGAAAAAATGGTCTCAAAGGGTGAAGAAGATAAC | Forward primer for replacing GFP with mCherry in pBAB7020 |
| | AA209-CAGTTGGAATTCTACGAATGGAATTCCTACTTATACAATTCATCCATGCC | Reverse primer for replacing GFP with mCherry in pBAB7020 |
| P*clc-3*::CLC-3(ΔPDZ)::T2A::mCherry | AA208-AGAATCCTGGCCCAGAAAAAATGGTCTCAAAGGGTGAAGAAGATAAC | Forward primer for replacing GFP with mCherry in pBAB7026 |
| | AA209-CAGTTGGAATTCTACGAATGGAATTCCTACTTATACAATTCATCCATGCC | Reverse primer for replacing GFP with mCherry in pBAB7026 |
| P*unc-17*::CLC-3(ΔPDZ)::T2A::mCherry | AA212-AATGAAATAAGCTTGCATGCGCACACACACACATCCTCTCTTC | Forward primer for replacing *clc-3* promoter with *unc-17* promoter in pBAB7027 |
| | AA213-CAAAGTGAACAAAACCCAACCTCTCTCTCTCCCCCTGGAATAT | Reverse primer for replacing *clc-3* promoter with *unc-17* promoter in pBAB7027 |
| P*unc-17*:: CLC-3::SpYFP(VC155) | AA218-ATCACTCTCGGCATGGACGAGCTGTACAAGTAACATTCGTAGAATTCCAACTGAGCGC | Forward primer for amplifying vector pBAB7025 |

| | | |
|---|---|---|
| | AA220-<br>TGTTTCAGATCGTTCGGAAT<br>TTTGCACGCCGGGCGGTAA<br>ACTTCAGTTACTGGCGGG | Reverse primer for amplifying vector pBAB7025 (overhang is composed of one half of the linker sequence separating CLC-3 and SpYFP) |
| | AA221-<br>ATTCCGAACGATCTGAAACA<br>GAAAGTGATGAACCATGAC<br>AAGCAGAAGAACGGCATCA | Forward primer for amplification of VC155 fragment of YFP (overhang is composed of second half of the linker sequence separating CLC-3 and SpYFP)<br><br>Linker sequence - cgcccggcgtgcaaaattccgaacgatctgaaacagaaagtgatgaaccat |
| | AA222-<br>CTCAGTTGGAATTCTACGAA<br>TGTTACTTGTACAGCTCGTC<br>CATGCC | Reverse primer for amplification of VC155 fragment of YFP |
| P*unc-17*::<br>CLC-3(ΔPDZ)::SpYFP(VC155) | AA218-<br>ATCACTCTCGGCATGGACG<br>AGCTGTACAAGTAACATTCG<br>TAGAATTCCAACTGAGCGC | Forward primer for amplifying vector pBAB7025 |
| | AA219-<br>TGTTTCAGATCGTTCGGAAT<br>TTTGCACGCCGGGCGTTCA<br>TATTCGACATCCGGGTCC | Reverse primer for amplifying vector pBAB7025 (overhang is composed of one half of the linker sequence separating CLC-3(ΔPDZ) and SpYFP) |
| | AA221-<br>ATTCCGAACGATCTGAAACA<br>GAAAGTGATGAACCATGAC<br>AAGCAGAAGAACGGCATCA | Forward primer for amplification of VC155 fragment of YFP (overhang is composed of second half of the linker sequence separating CLC-3(ΔPDZ) and SpYFP)<br><br>Linker sequence - cgcccggcgtgcaaaattccgaac |

| | | |
|---|---|---|
| | | gatctgaaacagaaagtgatgaac cat |
| | AA222- CTCAGTTGGAATTCTACGAA TGTTACTTGTACAGCTCGTC CATGCC | Reverse primer for amplification of VC155 fragment of YFP |
| P*unc-17*:: NAB-1::SpYFP(VN173) | AA218- ATCACTCTCGGCATGGACG AGCTGTACAAGTAACATTCG TAGAATTCCAACTGAGCGC | Forward primer for amplifying vector pBAB7032 |
| | AA224- TGTTTCAGATCGTTCGGAAT TTTGCACGCCGGGCGCATG GGAATTGTGTGTGCAATGA C | Reverse primer for amplifying vector pBAB7032 (overhang is composed of one half of the linker sequence separating NAB-1 and SpYFP) |
| | AA225- ATTCCGAACGATCTGAAACA GAAAGTGATGAACCATGTG AGCAAGGGCGAGGA | Forward primer for amplification of VN173 fragment of YFP (overhang is composed of second half of the linker sequence separating NAB-1 and SpYFP)<br><br>Linker sequence - cgcccggcgtgcaaaattccgaac gatctgaaacagaaagtgatgaac cat |

| | | |
|---|---|---|
| | AA226-CTCAGTTGGAATTCTACGAATGCTACTCGATGTTGTGGCGGA | Reverse primer for amplification of VN173 fragment of YFP |
| P*unc-17*::NAB-1::T2A::mCherry | AA214-GTACCGGTAGAGGGCAGAGGAAG | Forward primer for amplifying vector pBAB7025 |
| | AA215-GGTACCCTTGTTTTAGTAGGTTACTATTTTG | Reverse primer for amplifying vector pBAB7025 |
| | AA216-ACCTACTAAAACAAGGGTACCATGACAACGGCTTCCGAGCT | Forward primer for amplifying NAB-1 cDNA |
| | AA217-TCTGCCCTCTACCGGTACCATGGGAATTGTGTGTGCAATGA | Reverse primer for amplifying NAB-1 cDNA |

**Figure S1: Expression patterns of select claudin-like genes recapitulate previously described profiles**

(A) Transcriptional reporters for *hic-1*, *hpo-30*, *nsy-4*, and *vab-9* largely reproduce known expression patterns, with *vab-9* showing additional expression in tail neurons. Expression of *clc-11* transcript is limited to non-neuronal tissues. Scale bar = 100 mm

(B) The expression of *clc-9* transcript is localized to the intestine. Scale bar = 50 mm

**Figure S2: Mutants in *clc-3* do not show defects in locomotory speed**

(A) Schematic of the locomotion assay using the Worm Tracker system to record *C. elegans* movement and quantify parameters such as mean body-bend amplitude and speed.

(B) Representative tracks showing *C. elegans* movement during locomotion. Scale bar = 2 mm

(C) The speed of movement in *clc-3* mutants is comparable to that of WT *C. elegans*. *N*=3; *n*=10. Data are mean ± SD. Statistical analyses were performed using unpaired Student's *t* test.

(D) Time-course of aldicarb-induced paralysis. *clc-3* mutants exhibit accelerated paralysis compared to WT *C. elegans*. This phenotype is rescued in lines expressing CLC-3 under its endogenous or cholinergic neuron-specific promoter, with paralysis curves resembling those of WT animals. *N*≥3 ; *n*=20. Data are mean ± SEM

**Figure S3: Postsynaptic receptor sensitivity is unaltered in *clc-3* mutants.**

(A) Representative trace of muscle response to exogenously applied ACh. The amplitude of the response in *clc-3* mutants is comparable to WT animals, indicating that postsynaptic receptor sensitivity in these mutants is not compromised.

Numbers at the base of the bars indicate the number of *C. elegans* used for recordings. Data are mean ± SEM.

**Figure S4: Neuron and muscle development are unaffected in *clc-3* mutants**

(A) The density of SNB-1 puncta at cholinergic synapses in the dorsal nerve cord is unaltered in *clc-3* mutants. Numbers at the base of the bars indicate the number of *C. elegans* used for imaging. Data are mean ± SEM. Statistical analyses were performed using one-way ANOVA followed by Tukey's post-hoc test.

(B) Representative images showing cholinergic neuron architecture in WT and *clc-3* mutants. No defects were observed in the development or morphology of cholinergic neurons in *clc-3* mutants. Scale bar = 50 mm; *n*=10-15

(C) Representative images showing NMJ morphology marked by P*him-4*::MB::YFP in WT and *clc-3* mutant animals. NMJ morphology appears normal in *clc-3* mutants. Scale bar = 20 mm; *n*=10-15

(D) Representative images showing muscle structure as marked by MYO-3::GFP in WT and *clc-3* mutant animals. Myosin organization is normal in *clc-3* mutants. Scale bar = 20 mm; *n*=10-15

**Figure S5:** The density of SNB-1 puncta is unaffected across genotypes

(A) Time course of aldicarb-induced paralysis in rescue lines expressing CLC-3 lacking the PDZ bm under the endogenous promoter. The paralysis profile resembles that of *clc-3* mutants, indicating that the absence of PDZ bm prevents the rescue of the aldicarb hypersensitivity phenotype. Data are mean ± SEM.

(B) Time course of aldicarb-induced paralysis in rescue lines expressing CLC-3 lacking the PDZ bm under a cholinergic neuron-specific promoter. Similar to (B), the paralysis profile mirrors that of *clc-3* mutants, indicating that the PDZ bm is essential for rescuing aldicarb hypersensitivity.

(C) The density of SNB-1 puncta at cholinergic synapses along the dorsal nerve cord is comparable across WT *C. elegans*, *clc-3* mutants, and rescue lines expressing either full-length CLC-3 or CLC-3 lacking PDZ bm under a cholinergic promoter. Numbers at the base of the bars indicate the number of *C. elegans* used for imaging. Data are mean ± SEM. Statistical analyses were performed using one-way ANOVA followed by Tukey's post-hoc test.

**Figure S6: Spontaneous mEPSCs and mIPSCs are unaffected in *nab-1* and *nab-1; clc-3* double mutants**

(A) Time course of aldicarb-induced paralysis. *nab-1; clc-3* double mutants are hypersensitive to aldicarb, and their paralysis profile closely resembles that of *nab-1* and *clc-3* single mutants. *N*=3; *n*=20. Data are mean ± SEM.

(B) Representative traces of whole-cell recordings from body wall muscles showing miniature excitatory postsynaptic currents (mEPSCs). Quantification of mEPSC frequency and amplitude, shown in the accompanying bar graphs, indicates that these parameters are similar across WT, *clc-3* and *nab-1* single mutants, and *nab-1; clc-3* double mutant animals.

(C) Representative traces showing miniature inhibitory postsynaptic currents (mIPSC). Bar graphs indicate that mIPSC frequency and amplitude are similar across all genotypes.

Numbers at the base of the bars in (B) and (C) indicate the number of *C. elegans* used for recordings. Data are mean ± SEM.

(D) Bar graph quantifying the density of SNB-1 puncta at cholinergic synapses along the dorsal nerve cord. The number of puncta per micron is comparable across WT *C. elegans*, *nab-1; clc-3* double mutants and respective single mutants. Numbers at the base of the bars indicate the number of *C. elegans* used for imaging. Data are mean ± SEM. Statistical analyses were performed using one-way ANOVA followed by Tukey's post-hoc test.

**Figure S7: CLC-3 and NAB-1 physically interact with each other via the CLC-3 PDZ-binding motif in head cholinergic neurons**

(A) Representative images of the head region of WT animals expressing either CLC-3::SpYFP and NAB-1::SpYFP together, CLC-3(ΔPDZ)::SpYFP and NAB-1::SpYFP together, or only NAB-1::SpYFP in the cholinergic neurons. Clear YFP fluorescence can be seen in head cholinergic neurons when full-length CLC-3 and NAB-1 are co-expressed, but not in the other two conditions. Scale bar = 20 mm, *n*=10-17.

### Supplementary Figure 1

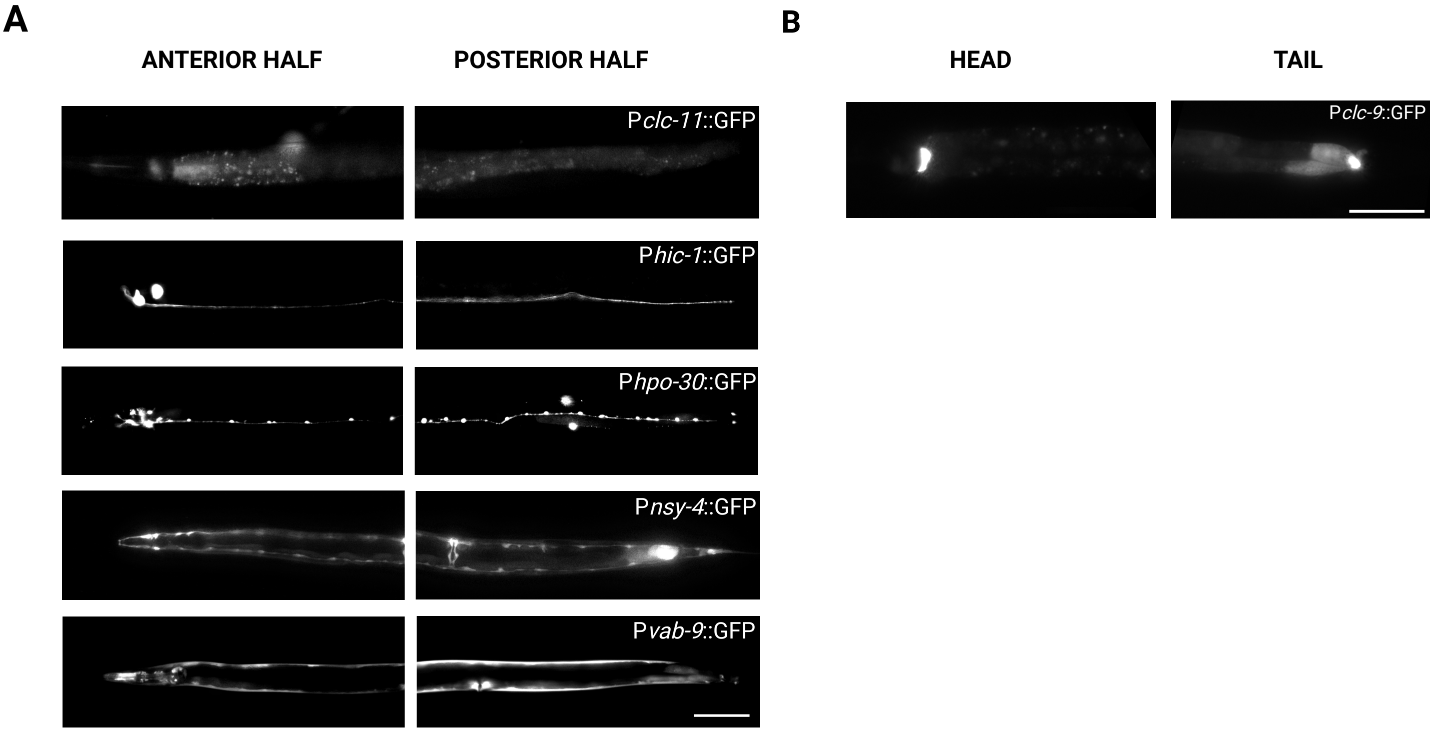

### Supplementary Figure 2

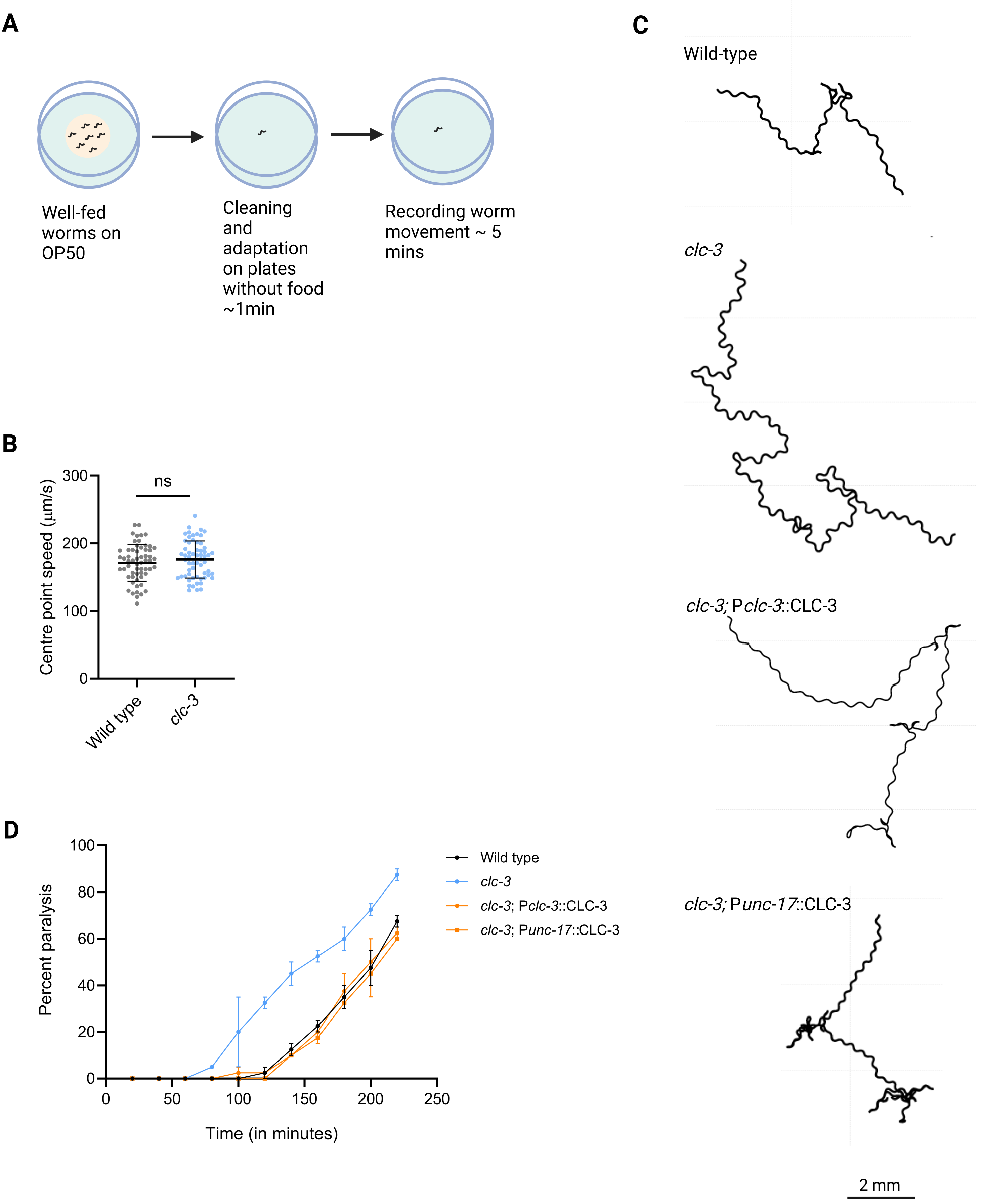

#### Supplementary Figure 3

**A**

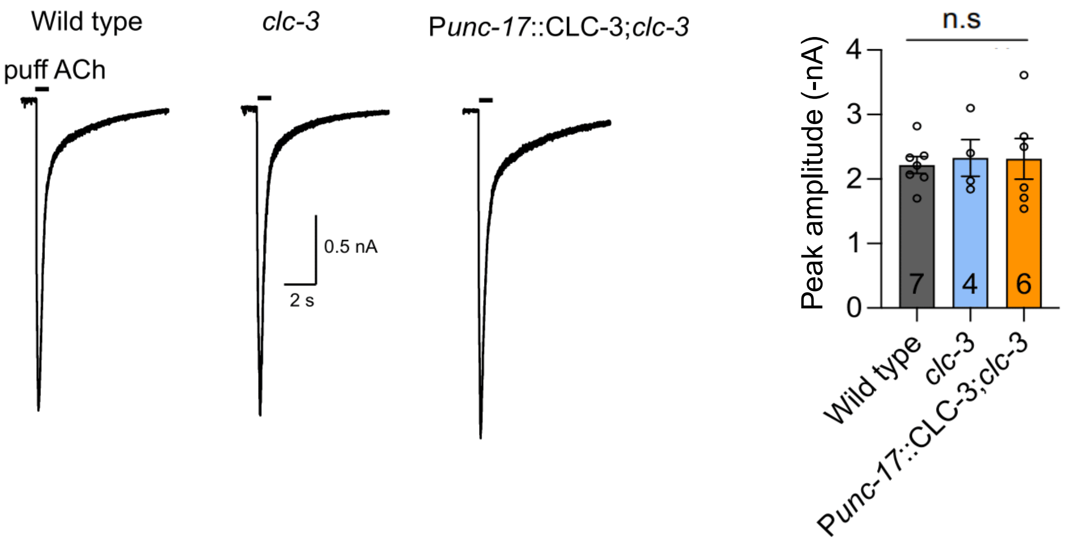

**Supplementary Figure 4**

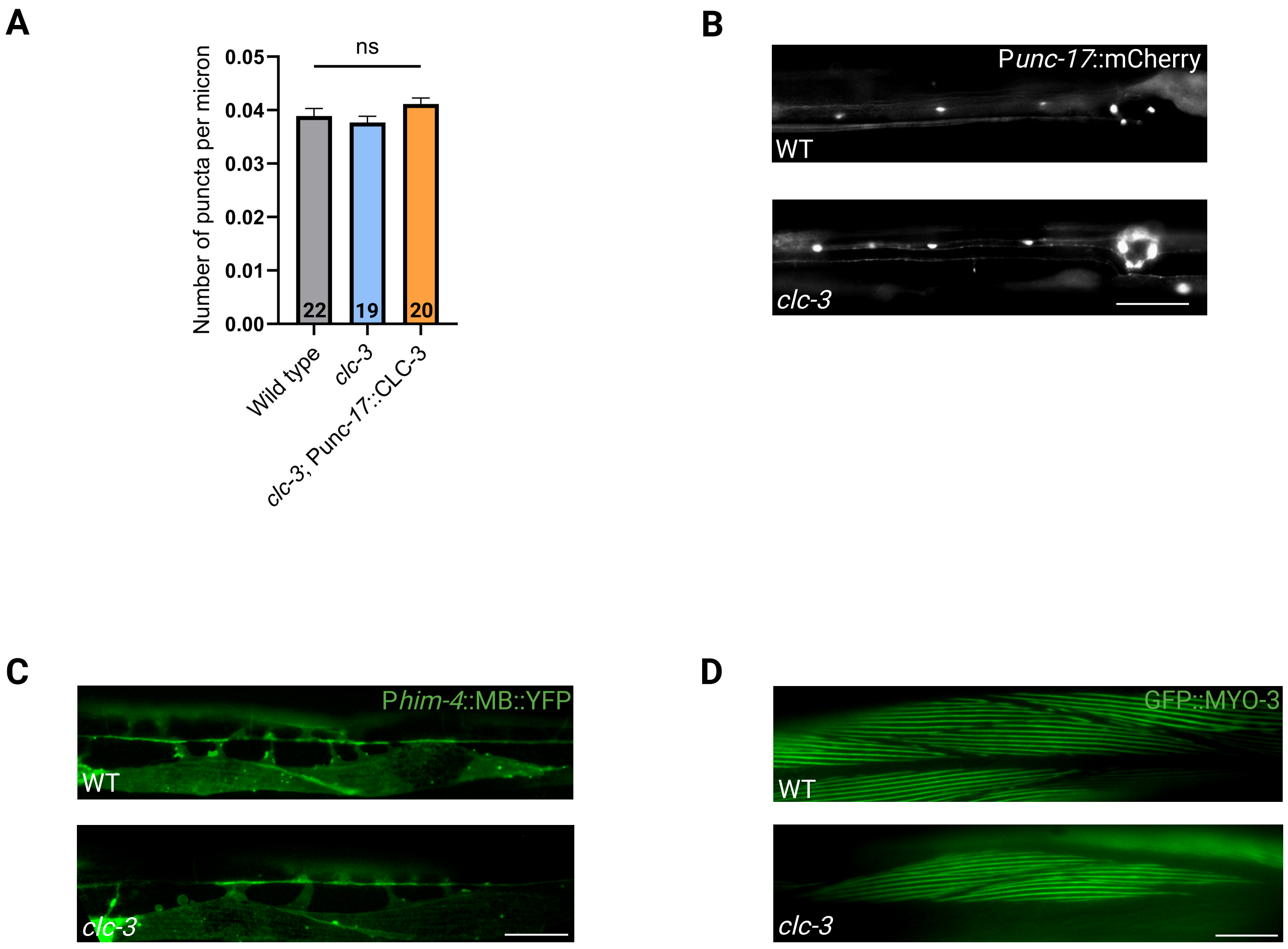

### Supplementary Figure 5

**A**

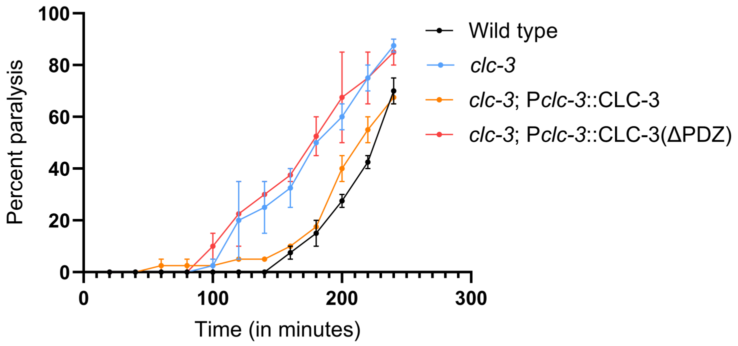

**B**

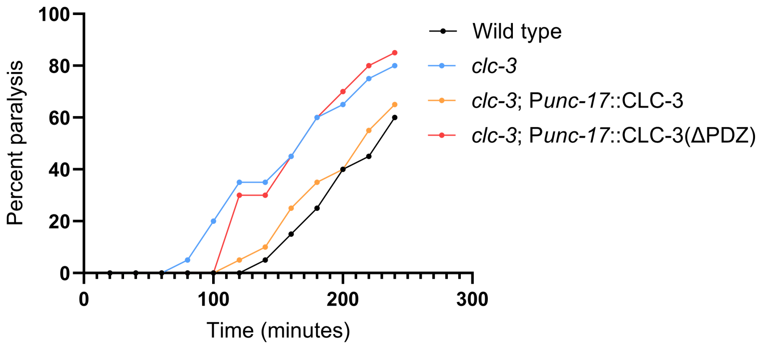

**C**

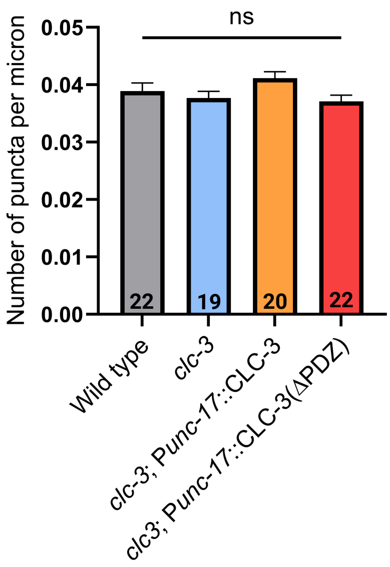

#### Supplementary Figure 6

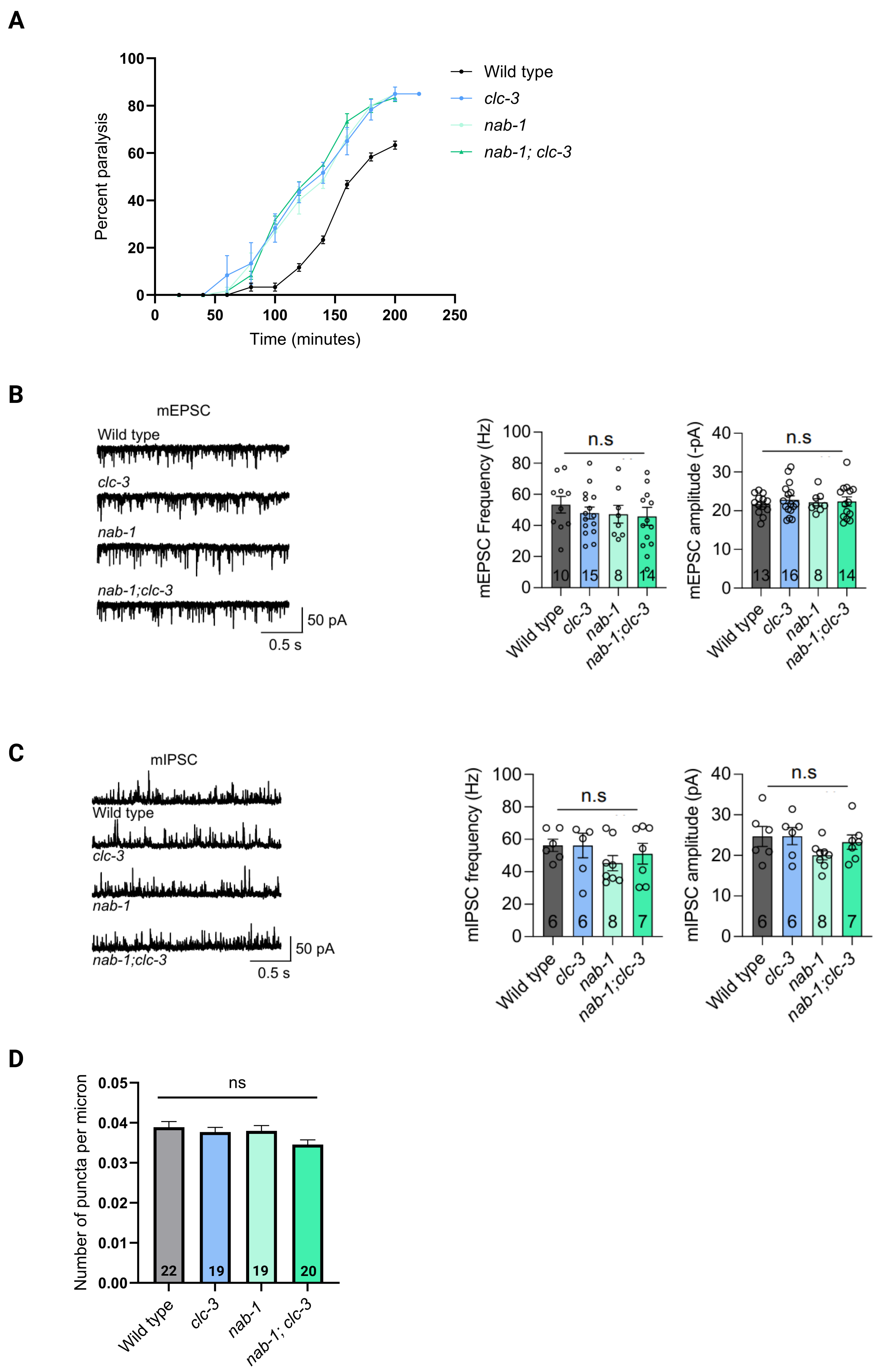

#### Supplementary Figure 7

**A**

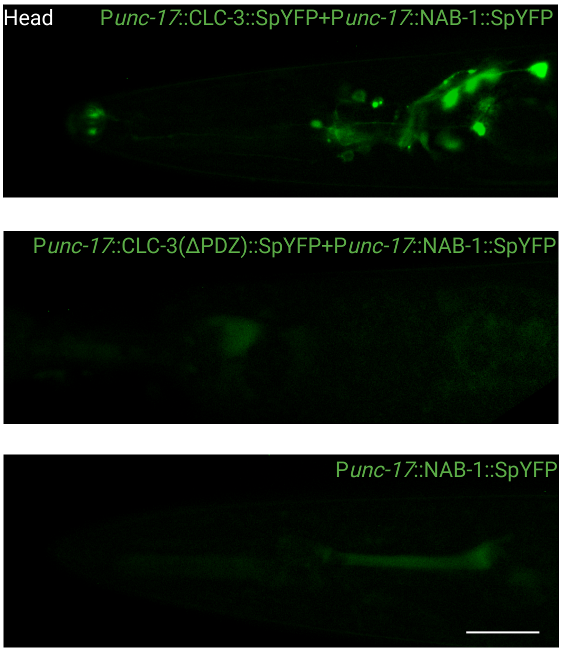
